# Atypical PI3Ks coordinate chemotaxis, signaling dynamics, and multicellular development in *Dictyostelium*

**DOI:** 10.64898/2026.05.11.724005

**Authors:** Laith Bahlouli, Erik Zhang, Eric Jung, Taylor Brentjens, Elizabeth Rose, Harrison Drebin, Ben Edwards, Isabella Wischik, Alyssa Huynh, Jacqueline Shen, Will Callahan, Kevin Zhangxu, Marc Edwards

## Abstract

Phosphoinositide 3-kinase (PI3K) signaling regulates protrusion, polarity, membrane uptake, and multicellular development in *Dictyostelium discoideum*, but these functions have been interpreted largely through canonical Class I PI3Ks and PI(3,4,5)P₃ production. This framework does not fully explain how PI3K-dependent pathways attenuate Ras activity, organize PI(3,4)P₂-associated polarity states, support cAMP relay, or coordinate development. Here, we identify three atypical PI3K-family enzymes—PikF, PikG, and PikH—as functionally distinct regulators of these processes. PikF constrains Ras–phosphoinositide–actin signaling; pikF⁻ cells show prolonged cAMP-stimulated Ras activation, extended PIP₃ recruitment, delayed PI(3,4)P₂ biosensor recovery, elevated peripheral actin activity, impaired chemotactic precision, and delayed abnormal development. PikG acts through a distinct relay-associated pathway: pikG⁻ cells fail to generate endogenous cAMP oscillations, display disrupted ACA polarity, deposit spatially disorganized ACA-positive vesicle trails, and fail to aggregate. PikH, in contrast, supports efficient phagocytic uptake with little effect on acute chemotactic signaling. Kinase-dead rescue experiments show that conserved catalytic lysines are required for PikF- and PikG-dependent development and PikH-dependent uptake. Together, our results reveal that atypical PI3Ks diversify the *Dictyostelium* PI3K signaling toolkit, separating protrusive signal attenuation, cAMP relay organization, membrane uptake, and multicellular development into distinct kinase-dependent modules.

## Introduction

Cell migration underlies a wide range of physiological and pathological processes, including multicellular development, immune surveillance, wound healing, and cancer metastasis ((Ridley *et al*., 2003; Garcia and Parent, 2008; Friedl and Wolf, 2010). Directed migration depends on coordinated signaling and cytoskeletal remodeling, which together generate polarized cell states characterized by spatially confined protrusive activity at the front and contractile activity at the rear (Swaney *et al*., 2015; Yamada and Sixt, 2019). In amoeboid cells ranging from *Dictyostelium discoideum* to neutrophils, front–back domains can emerge even in the absence of external gradients, consistent with models in which polarity is governed by excitable signal-transduction networks shaped by coupled positive- and negative-feedback loops (Devreotes *et al*., 2017; Pal *et al*., 2019). In *Dictyostelium*, starvation-induced development uses these polarity and excitability networks to build a multicellular structure from a field of individual cells (Loomis, 2014). Periodic, cyclic adenosine monophosphate (cAMP) signaling provides both a communication system between cells and a navigational cue that engages the polarity machinery within each cell (Hashimura *et al*., 2019). In response, cells polarize, migrate, and stream toward aggregation centers, forming multicellular mounds that undergo morphogenesis to generate fruiting bodies (Dormann and Weijer, 2003; Kimmel and Parent, 2003; Weijer, 2004).

Phosphoinositide signaling is a central component of amoeboid polarity networks. In *Dictyostelium* and neutrophil chemotaxis, Ras activation, PI3K activity, and PI(3,4,5)P₃ accumulation are associated with front-state signaling, whereas PTEN, myosin II, and PI(3,4)P₂-enriched domains are associated with complementary rear-state organization (Charest and Firtel, 2006; Swaney, Huang and Devreotes, 2010; Lam *et al*., 2012; Li *et al*., 2021). Together, these opposing front- and rear-state modules form a partially redundant signaling system that regulates protrusion dynamics, inhibitory control, and actin organization. Although PI(3,4,5)P₃ is enriched at the front of cells during chemotaxis, work in *Dictyostelium* and neutrophils has shown that PI3K/PIP₃ signaling is not the sole determinant of directional sensing or migration (Ferguson *et al*., 2007; Hoeller and Kay, 2007). Instead, PI3K/PIP₃ signaling contributes to the biasing, amplification, and persistence of protrusive activity, shaping how pseudopods are generated, split, selected, and maintained during migration (Postma *et al*., 2004; Andrew and Insall, 2007). These same properties become central during development, when protrusive behavior and polarity must be coordinated for cAMP-mediated aggregation (Dormann *et al*., 2002; Weijer, 2004).

Despite this broad functional scope, *Dictyostelium* PI3K signaling has largely been interpreted through the lens of canonical PI(3,4,5)P₃-generating Class I enzymes. This framework is grounded in genetic studies linking *Dictyostelium* Class I PI3Ks to chemoattractant-stimulated PI(3,4,5)P₃ production, chemotactic signaling, and PI3K-dependent macropinocytosis (Funamoto *et al*., 2002; Hoeller and Kay, 2007; Hoeller *et al*., 2013). Similarly, perturbing PI(3,4,5)P₃ turnover through loss of the phosphatases PTEN and Dd5P4 disrupts the timing and spatial organization of phosphoinositide signaling across processes ranging from migration to phagocytosis and macropinocytosis (Iijima and Devreotes, 2002; Loovers, 2007). However, several observations remain difficult to explain from a Class I-centered framework alone. These include the timing of protrusive signaling during Ras/PI3K excitability, PI3K-dependent control of Adenylyl Cyclase A, ACA, activation and adaptation, and the organization of PI(3,4)P₂-enriched rear and endocytic membrane domains (Postma *et al*., 2004; Comer and Parent, 2006; Li *et al*., 2018, 2021).

The source and spatial organization of PI(3,4)P₂ are especially difficult to reconcile with a strictly Class I PI3K-centered model. Although PI(3,4)P₂ has often been viewed as a product of PI(3,4,5)P₃ turnover by inositol 5-phosphatases, including SHIP-family enzymes in mammalian cells and the OCRL homolog Dd5P4 in *Dictyostelium*, it has emerged as a spatially organized signaling lipid in its own right (Loovers, 2007; Nishio, 2007; Phillip T. Hawkins and Stephens, 2016). (PI(3,4)P₂ has been implicated in endocytic and phagocytic membrane remodeling, cytoskeletal regulation, Ras-dependent polarity, and rear-state identity (Loovers, 2007; Posor *et al*., 2013; P. T. Hawkins and Stephens, 2016; Li *et al*., 2021). Recent work in *Dictyostelium* showed that PI(3,4)P₂ persists at the plasma membrane after pharmacological inhibition of PI3K-driven PI(3,4,5)P₃ production and continues to accumulate at rear membranes and macropinosomes in PI3KA/B-null cells (Li *et al*., 2021). These observations suggest that a population of PI(3,4)P₂ is generated independently of canonical Class I PI3K-mediated PI(3,4,5)P₃ production, potentially through alternative phosphoinositide-conversion routes involving PI4P and/or PI3P (Posor *et al*., 2013; P. T. Hawkins and Stephens, 2016; Wallroth and Haucke, 2018; Li *et al*., 2021).

The persistence of PI(3,4)P₂ outside canonical Class I PI(3,4,5)P₃ production shifts attention to non-Class I PI3K-family enzymes as potential contributors to *Dictyostelium* polarity and development. Class III PI3Ks generate PI3P, whereas metazoan Class II PI3Ks can generate PI(3,4)P₂ from PI4P and regulate endocytic membrane remodeling (Balla, 2013; Posor *et al*., 2013; P. T. Hawkins and Stephens, 2016; Wallroth and Haucke, 2018; Gulluni *et al*., 2019). In *Dictyostelium*, the Class III branch is represented by the Vps34-like kinase PikE/DdPIK5, which has been linked to protein trafficking, growth, and multicellular development (Zhou *et al*., 1995). However, *Dictyostelium* appears to lack canonical Class II PI3Ks (Engelman, Luo and Cantley, 2006; Brown and Auger, 2011) creating an unresolved gap between PI(3,4)P₂-dependent behaviors and the enzymes currently known to generate or remodel this lipid. This gap motivates a broader question: whether additional, atypical PI3K-family enzymes help account for PI3K-dependent control of protrusion, uptake, signaling dynamics, and developmental coordination that cannot be fully explained by the canonical Class I PI(3,4,5)P₃ pathway.

Here we identify and characterize three atypical PI3Ks—PikF, PikG, and PikH—as regulators of PI3K-dependent behavior in *Dictyostelium*. Comparative sequence analysis and phylogenetic placement suggest that these enzymes diverge from canonical Class I PI3Ks at activation-loop features predicted to influence phosphoinositide substrate engagement(Pirola *et al*., 2001).

Functionally, these enzymes define distinct, nonredundant PI3K-dependent modules: PikH supports efficient phagocytic uptake, PikF limits the duration and spatial organization of protrusive signaling, and PikG is required for endogenous cAMP oscillations and aggregation-stage coordination. Together, these findings expand the landscape of PI3K signaling in *Dictyostelium* and identify atypical PI3Ks as regulators of membrane uptake, chemotaxis, and developmental coordination beyond the canonical Class I framework.

## Results

### PikF, PikG, and PikH are atypical PI3K-family enzymes distinct from canonical Class I PI3Ks

To determine how PI3K signaling is diversified beyond canonical Class I output in *Dictyostelium*, we analyzed PikF, PikG, and PikH using comparative sequence analysis and cell biological assays. Alignment of the PI3K activation-loop region showed that PikF, PikG, and PikH lack the positively charged residue predicted to stabilize the 5′ phosphate of PI(4,5)P_2_, distinguishing them from canonical Class I PI3Ks in both humans and *Dictyostelium* (Fig. 1A,B). Phylogenetic analysis similarly separated PikF and PikG from the Class I-like PikA/PikB/PikC clade, while PikH clustered closer to Vps34/Class III-like PI3Ks (Fig. 1C). Comparative sequence analysis suggested that PikF, PikG, and PikH represent atypical PI3Ks with signaling outputs distinct from canonical Class I enzymes.PikF and PikG regulate vegetative motility, but PikF uniquely disrupts chemotactic directionality

**Figure 1.**
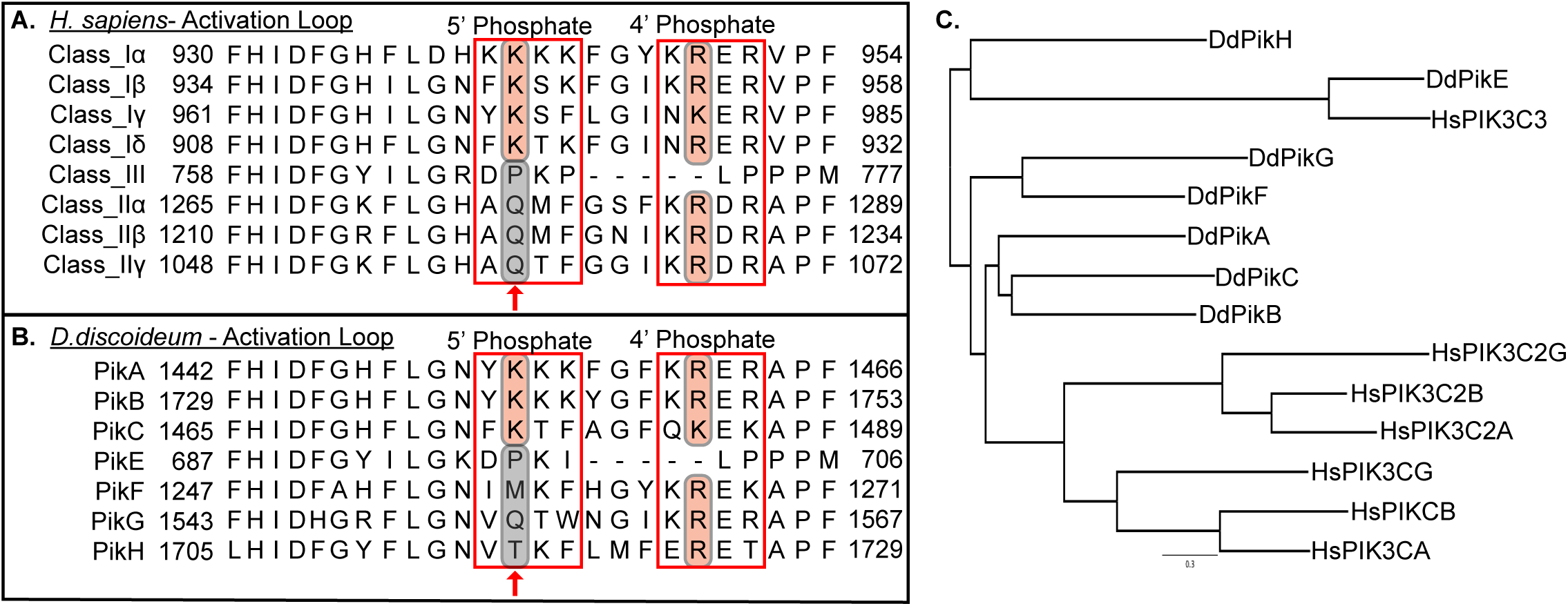
Activation-loop sequence alignment and guide-tree comparison of human and *Dictyostelium* PI3Ks. **(A,B)** Alignment of activation-loop sequences from human **(A)** and *Dictyostelium discoideum* (B) PI3Ks. Residues in the putative 4′- and 5′-phosphate–stabilizing basic boxes are highlighted in red. Arrows indicate the conserved lysine present in Class I PI3Ks but absent from human Class II and Class III PI3Ks, *Dictyostelium* PikE, and PikF–H. **(C)** Maximum-likelihood guide tree generated from human and *D. discoideum* PI3K activation-loop sequences. The scale bar indicates evolutionary distance. PikF, PikG, and PikH group separately from the canonical *Dictyostelium* Class I PI3Ks PikA, PikB, and PikC, consistent with divergence at activation-loop features that distinguish Class I from non-Class I PI3K branches.

### PikF and PikG regulate vegetative motility, but PikF uniquely disrupts chemotactic directionality

PI3Ks regulate multiple actin-dependent processes, so we first tested whether PikF, PikG, and PikH contribute to motility in vegetative cells. We performed time-lapse motility experiments by widefield imaging vegetative-stage Ax2 wild-type, pikF⁻, pikG⁻, and pikH⁻ cells for 1 h, followed by single-cell tracking. pikH⁻ cells behaved most similarly to Ax2 cells, whereas *pikF⁻* and *pikG⁻* cells exhibited less persistent trajectories (Fig. 2A–D). Quantification confirmed pronounced reductions in both speed and persistence in *pikF⁻* and *pikG⁻* cells, with median values reduced by approximately 40% for speed and 50% for persistence relative to Ax2 in both mutants (Fig. 2E,F). Kruskal–Wallis analysis confirmed significant reductions in speed and persistence for *pikF⁻* and *pikG⁻* cells, whereas *pikH⁻* cells did not differ significantly from Ax2.

**Fig. 2.**
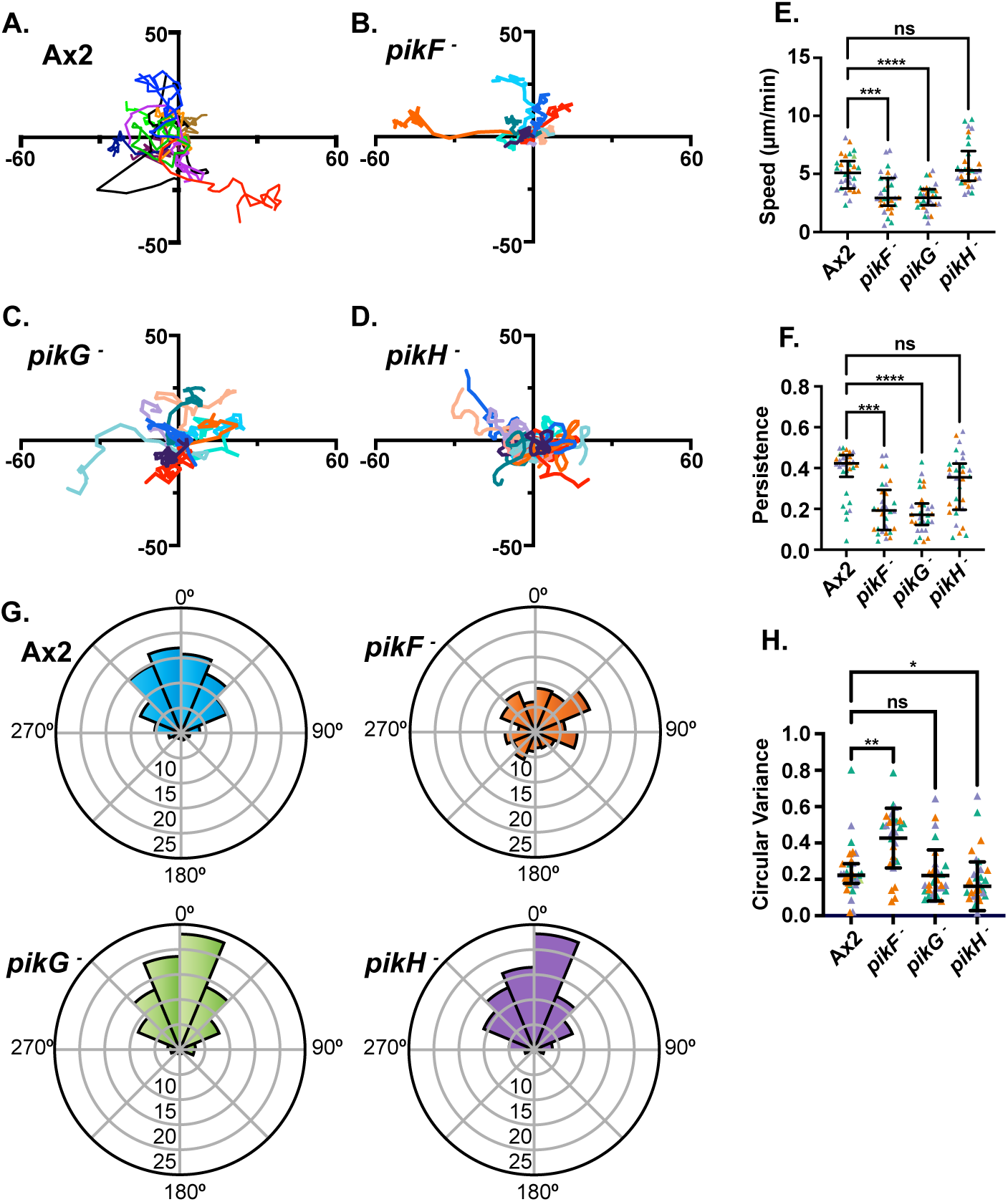
Atypical PI3Ks differentially regulate random migration and chemotactic accuracy in *Dictyostelium*. **(A–D)** Random-migration tracks of representative Ax2, *pikF*⁻, *pikG⁻*, and *pikH⁻* cells. Ten cells per strain were tracked at 2-min intervals for 60 min and plotted from a common origin; axes are in µm. **(E)** Cell speed during random migration. Each point represents one cell; n = 30 cells per strain from three biological replicates, with 10 cells analyzed per replicate. Replicates are distinguished by color. Bars indicate median and interquartile range (IQR). **(F)** Persistence of cells tracked as in E. n = 30 cells per strain from three biological replicates. Replicates are distinguished by color. Bars indicate median and IQR. **(G)** Rose plots showing the protrusion angle θ between each cell step and the direction of an ideal path toward the cAMP-filled microcapillary. Bars show the proportion of steps in each angular bin. For each strain, 100 cells across three biological replicates were analyzed, with 100 steps analyzed per cell. **(H)** Circular variance for the indicated strains. n = 30 cells per strain from three biological replicates. Replicates are distinguished by color. Bars indicate median and IQR. For E, F, and H, significance was assessed by Kruskal–Wallis test with Dunn’s multiple-comparisons test. Asterisks indicate significant differences (P < 0.001); ns, not significant.

We then asked whether these motility defects also altered chemotaxis toward an exogenous cAMP signal. Cells were exposed to a microcapillary containing 10 μM cAMP and imaged by time-lapse microscopy. To quantify chemotactic accuracy, pseudopod angles were measured relative to a hypothetical straight-line path toward the needle, with smaller angles indicating closer alignment to the gradient. These angles were then plotted as angular distributions for each strain (Fig. 2G). Ax2 cells showed a strong bias toward the cAMP source, with pseudopods tightly clustered within 45° of the needle axis (Fig. 2G; Fig. S1A; Movie 1). By contrast, *pikF⁻* cells displayed broader pseudopod-angle distributions and more meandering trajectories toward the needle, often through wide-angle lateral steps (Fig. 2G; Fig. S1B; Movie 1). This defect paralleled the reduced persistence observed during vegetative motility (Fig. 2B,F). *pikG⁻* and *pikH⁻* cells showed angular distributions more similar to Ax2 (Fig. 2G; Movies 2, 3), despite the reduced speed and persistence observed for *pikG⁻* cells during random migration (Fig. 2C, E, F). Quantification of circular variance confirmed a significant increase in directional dispersion of pseudopod angles in *pikF⁻* cells, but not in *pikG⁻* or *pikH⁻* cells, relative to Ax2 (Fig. 2H).

### PikF and PikG produce separable defects in aggregation and morphogenesis

Chemotaxis drives aggregation-stage streaming and multicellular development, raising the possibility that the migration defects in *pikF⁻* and *pikG⁻* cells alter developmental progression. We examined development on non-nutrient agar over 48 h using brightfield time-lapse imaging (Fig. 3A; Movie 4). *pikH⁻* cells developed similarly to Ax2, forming mounds, slugs, and fruiting bodies at approximately 8, 16, and 24 h, respectively, consistent with previous reports (Weijer, 2004) (Fig. 3A–C; Fig. S2A; Movie 4).

**Fig. 3.**
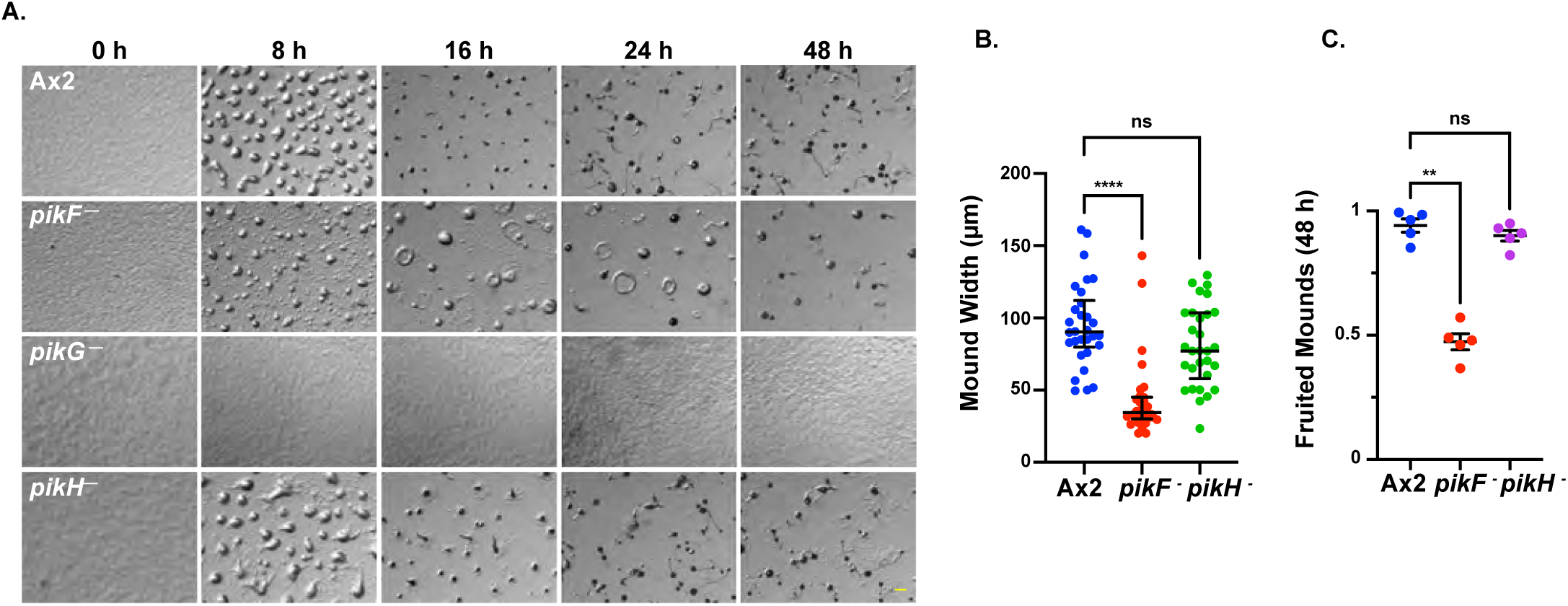
PikF and PikG differentially regulate multicellular development in *Dictyostelium*. **(A)** Representative time course of development on non-nutrient agar for Ax2, *pikF⁻*, *pikG⁻*, and *pikH⁻* cells at the indicated times. Images were acquired over 48 h at 7.3× magnification. n = 6 independent experiments. Scale bar, 500 μm. **(B)** Quantification of mound width from the experiments shown in A. n = 30 mounds per strain from three biological replicates. Bars indicate median and IQR. **(C)** Fraction of mounds reaching the fruiting-body stage by 48 h. Five biological replicates were analyzed, with approximately 30 mounds scored per plate. Bars indicate mean ± s.e.m. *pikF⁻* cells showed delayed development with abnormal mounds and slugs, whereas *pikG⁻* cells failed to complete development. *pikH⁻* development was similar to Ax2. Statistical significance was assessed by Kruskal-Wallis test; *P < 0.001 where indicated.

In contrast, *pikF⁻* cells showed delayed and abnormal development. Mound formation was delayed by 1–2 h relative to AX2, and the mounds that formed were significantly smaller (Fig. 3A,B; Fig. S2A; Movie 4). Subsequent morphogenesis was also disrupted: *pikF⁻* cells formed small rotating structures instead of normal slugs around 20 h, and fewer than 50% of mounds reached the fruiting-body stage by 48 h, whereas greater than 95% of AX2 aggregates did so by the same time point (Fig. 3A,C; Fig. S2B; Movie 4). *pikG⁻* cells showed the most severe developmental defect, failing to initiate streaming or aggregation even after 48 h (Fig. 3A; Fig. S2A; Movie 4). These data separate the delayed, abnormal morphogenesis of *pikF⁻* cells from the more severe aggregation-stage block in *pikG⁻* cells.

### PikF and PikH support actin-dependent membrane uptake

PI3Ks also regulate actin-dependent membrane remodeling, so we broadened our analysis to phagocytosis and macropinocytosis. *pikG⁻* cells behaved similarly to Ax2 across all uptake assays, whereas *pikF⁻* and *pikH⁻* cells showed reduced phagocytosis of fluorescent beads and zymosan particles. *pikF⁻* cells also showed reduced dextran uptake (Fig. 4A–C). Mixed-effects analysis of uptake kinetics confirmed significant defects in phagocytosis for *pikF⁻* and *pikH⁻* cells and in macropinocytosis for *pikF⁻* cells. Consistent with a kinase-dependent role for PikH, re-expression of wild-type PikH restored phagocytic uptake in *pikH⁻* cells, whereas kinase-dead PikH^K1586A failed to rescue the defect (Fig. S3). These uptake phenotypes broadened the physiological effects of PikF and PikH, while the strongest developmental phenotypes remained associated with *pikF⁻* and *pikG⁻* cells. We therefore focused next on cAMP-stimulated phosphoinositide signaling dynamics in all three mutants.

**Fig. 4.**
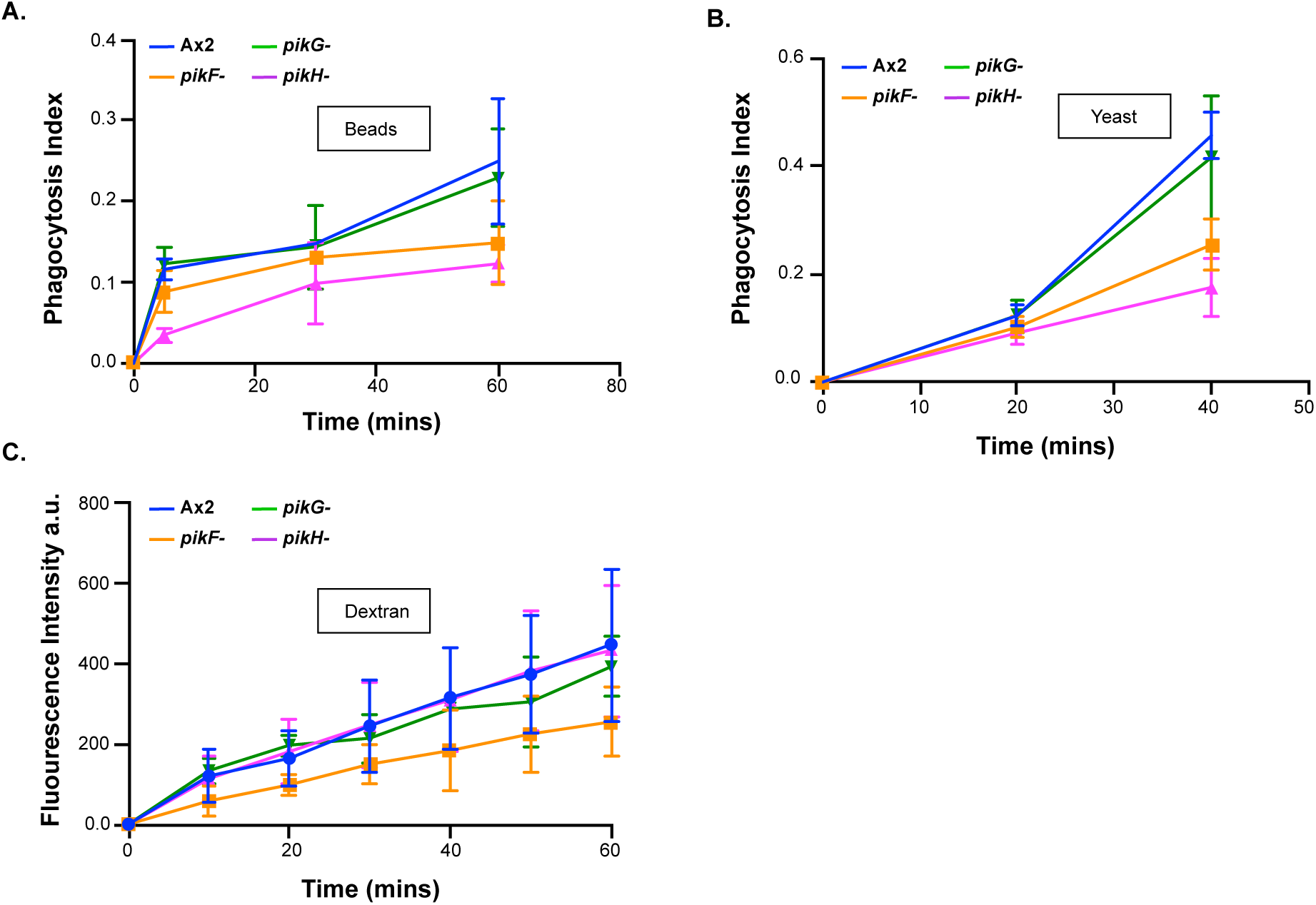
PikF and PikH differentially regulate uptake in *Dictyostelium*. **(A)** Phagocytosis index for uptake of fluorescent 4 μm beads by Ax2, *pikF⁻*, *pikG⁻*, and *pikH⁻* cells. Phagocytosis was quantified as the fraction of cells internalizing beads over time. n = 60 cells per strain from three biological replicates. **(B)** Phagocytosis index for uptake of fluorescent *Saccharomyces cerevisiae* particles by the indicated strains, quantified as in A. n = 60 cells per strain from three biological replicates. **(C)** Macropinocytosis measured by uptake of fluorescent dextran over time and quantified as the increase in intracellular fluorescence. n = 30 cells per strain from three biological replicates. Data are plotted as mean ± s.e.m. Statistical significance was assessed by mixed-effects analysis of uptake kinetics. *pikF⁻* and *pikH⁻* cells showed significant defects in phagocytosis (P < 0.0001), and *pikF⁻* cells showed a significant defect in macropinocytosis (P < 0.001)

### PikF prolongs cAMP-stimulated phosphoinositide signaling

The distinct effects of PikF, PikG, and PikH on migration, development, and uptake suggested that these kinases regulate actin-dependent behaviors through different signaling mechanisms. Because PI3Ks generate 3-phosphorylated phosphoinositides, we next asked whether the phenotypic differences among *pikF⁻*, *pikG⁻*, and *pikH⁻* cells were accompanied by differences in PI(3,4,5)P₃ and PI(3,4)P₂ dynamics. Cells expressing PH-CRAC-GFP or tPH-CynA-GFP were treated with latrunculin and stimulated uniformly with 10 μM cAMP to monitor PI(3,4,5)P₃ and PI(3,4)P₂ biosensor recruitment, respectively (Fig. 5). Latrunculin treatment suppressed actin-driven shape changes, allowing cAMP-dependent membrane recruitment to be quantified without changes in cell geometry confounding the analysis.

**Fig. 5.**
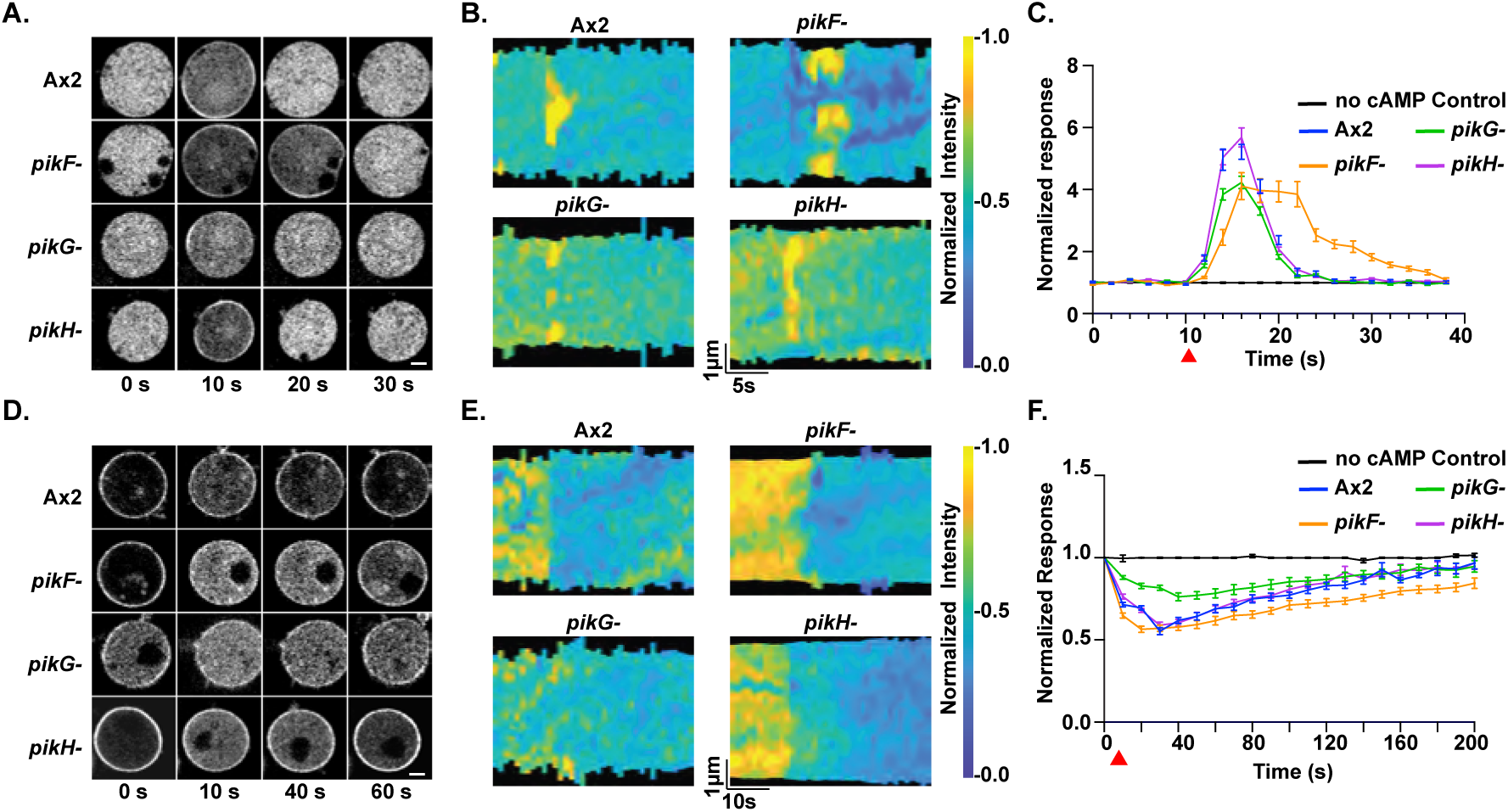
PikF prolongs phosphoinositide signaling dynamics, whereas PikG blunts acute responses to cAMP stimulation. **(A)** Representative time course of PH-CRAC-GFP, a PI(3,4,5)P₃ biosensor, in latrunculin-treated Ax2, *pikF⁻*, *pikG⁻*, and *pikH⁻* cells stimulated uniformly with 10 µM cAMP. Cells were imaged over 40 s. Scale bar, 2 µm. **(B)** Representative kymographs of the cells shown in A. **(C)** Baseline-normalized membrane-to-cytosol ratio of PH-CRAC-GFP following cAMP stimulation. cAMP was added at 10 s, indicated by the red arrow. n = 60 cells per strain from three biological replicates, with 20 cells analyzed per replicate. Data are plotted as mean ± s.e.m. **(D)** Representative time course of tPH-CynA-GFP, a PI(3,4)P₂ biosensor, in latrunculin-treated Ax2, *pikF⁻*, *pikG⁻*, and *pikH⁻* cells stimulated uniformly with 10 µM cAMP. Cells were imaged over 200 s. **(E)** Representative kymographs of the cells shown in D. **(F)** Baseline-normalized membrane-to-cytosol ratio of tPH-CynA-GFP following cAMP stimulation. cAMP was added at 10 s, indicated by the red arrow. n = 60 cells per strain from three biological replicates, with 20 cells analyzed per replicate. Data are plotted as mean ± s.e.m.

In Ax2 and *pikH⁻* cells, PH-CRAC-GFP rapidly accumulated at the membrane after cAMP stimulation, appearing in time-course images and kymographs as a brief membrane translocation followed by signal decay (Fig. 5A,B). Quantification of the membrane-to-cytosol ratio confirmed similar response and recovery kinetics in Ax2 and *pikH⁻* cells, with statistically indistinguishable peak responses and return to baseline within approximately 20–25 s of stimulation (Fig. 5C; Fig. S4A). By contrast, *pikF⁻* cells showed prolonged PH-CRAC-GFP membrane recruitment, with recovery times approximately 20% longer than Ax2 despite a similar peak response (P < 0.01; Fig. 5A–C; Fig. S4A). *pikG⁻* cells displayed response and recovery kinetics similar to Ax2, but with an approximately 20% reduction in peak response amplitude, consistent with a blunted acute PI(3,4,5)P₃ response after cAMP stimulation (Fig. 5A–C; Fig. S4A).

We observed a related pattern using tPH-CynA-GFP to monitor PI(3,4)P₂ dynamics. In Ax2 and *pikH*⁻ cells, cAMP stimulation caused a rapid decrease in membrane-associated tPH-CynA-GFP signal, visible in both the time-course images and kymographs, followed by gradual recovery toward baseline (Fig. 5D,E). Quantification of the membrane-to-cytosol ratio confirmed similar response and recovery kinetics in Ax2 and *pikH*⁻ cells (Fig. 5F; Fig. S4B). *pikF*⁻ cells showed a comparable initial decrease in tPH-CynA-GFP signal but significantly delayed recovery to 50% baseline, with recovery times approximately 25% longer than Ax2 (P<0.05; Fig. 5D–F; Fig. S4B). *pikG*⁻ cells recovered with kinetics similar to Ax2 but showed an approximately 20% smaller stimulation-induced decrease in membrane-to-cytosol ratio (P<0.05; Fig. 5D–F; Fig. S4B). Because *pikH*⁻ cells showed largely Ax2-like migration, development, and cAMP-stimulated phosphoinositide responses, subsequent mechanistic experiments focused on the more pronounced *pikF*⁻ and *pikG*⁻ phenotypes.Biochemical PIP3 measurements support a defect in signal attenuation in *pikF*⁻ cells Mass ELISA provided an independent biochemical measure of the PIP_3_ dynamics observed with PH-CRAC-GFP. Basal levels of PI(3,4,5)P_3_, PI(3,4)P_2_, and PI3P were not significantly different across Ax2, *pikF⁻*, *pikG⁻*, and *pikH⁻* cells, indicating that loss of these enzymes did not broadly alter steady-state levels of 3-phosphorylated phosphoinositides (Fig. S5A–C). We then measured cAMP-stimulated PI(3,4,5)P_3_ dynamics. In Ax2 cells, PI(3,4,5)P_3_ increased rapidly after cAMP stimulation, peaking approximately 5–8 s after stimulation and then declining back to baseline (Fig. S5D). *pikF⁻* cells showed a more sustained PI(3,4,5)P_3_ response, with an extended recovery phase that mirrored the prolonged PH-CRAC-GFP recruitment observed in live-cell biosensor assays (Fig. 5A–C; Fig. S4A; Fig. S5D). Mixed-effects analysis of the cAMP-stimulated time course identified a significant difference between Ax2 and *pikF⁻* cells over time. These biochemical measurements further support a specific defect in PI(3,4,5)P_3_ signal attenuation in *pikF⁻* cells.

### Loss of PikF increases peripheral actin activity and prolongs Ras activation

The prolonged PIP_3_ response in *pikF⁻* cells, together with wide-angle protrusive behavior during chemotaxis, suggested that loss of PikF is associated with altered downstream actin dynamics. Peripheral actin activity was assessed in Ax2 and *pikF⁻* cells expressing LimE-RFP, a marker of filamentous actin assembly. Cells were imaged for 500 s, and peripheral actin patches were quantified as described in the Methods (Fig. S6A,B). *pikF⁻* cells displayed significantly more peripheral actin patches per cell than Ax2 cells (, indicating elevated cortical actin activity (Fig. S6A). Representative kymographs showed the same pattern, with *pikF⁻* cells exhibiting more frequent peripheral actin patches over time than Ax2 cells (Fig. S6B).

The combination of prolonged phosphoinositide signaling and elevated peripheral actin activity pointed to Ras as a candidate contributor to the *pikF⁻* protrusive phenotype. Ras activation was monitored using the RBD-GFP biosensor in latrunculin-treated Ax2 and *pikF⁻* cells stimulated with uniform 10 μM cAMP (Fig. 6A,B). As with the PH-CRAC-GFP and tPH-CynA-GFP biosensor assays, RBD-GFP recruitment was quantified as a baseline-normalized membrane-to-cytosol ratio (Fig. 6B). In Ax2 cells, RBD-GFP rapidly accumulated at the membrane, peaked approximately 6 s after stimulation, and returned to baseline by approximately 20 s (Fig. 6A,B). In *pikF⁻* cells, RBD-GFP showed a similar initial response but remained elevated at 24 s after stimulation, indicating prolonged Ras activation (Fig. 6A,B).

**Fig. 6.**
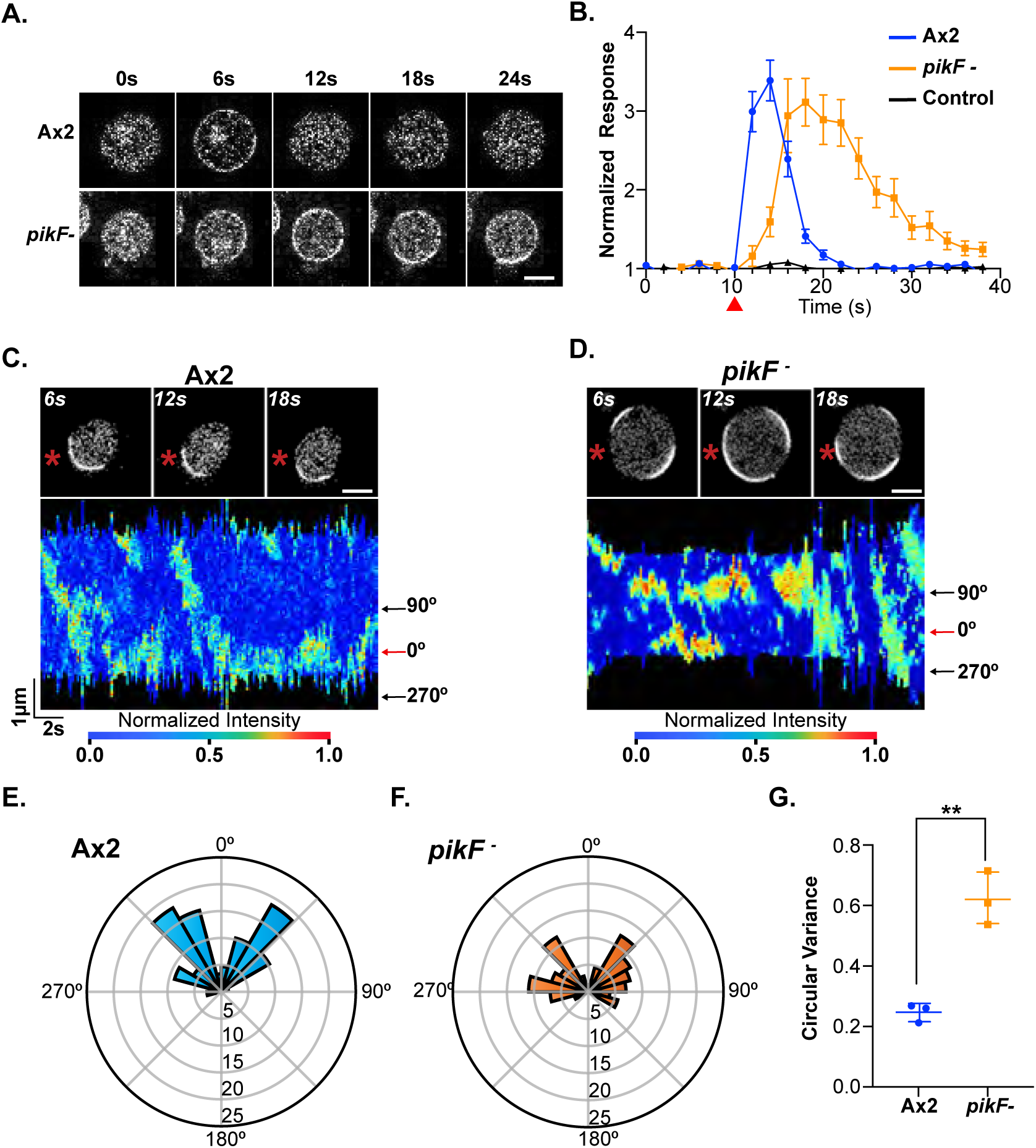
PikF prolongs Ras activation and impairs Ras crescent positioning. **(A)** Representative time course of Ras activation in latrunculin-treated Ax2 and *pikF⁻* cells expressing RBD-GFP following uniform stimulation with 10 µM cAMP. Cells were imaged every 2 s over 40 s; cAMP was added at 10 s. Scale bar, 10 µm. **(B)** Baseline-normalized membrane-to-cytosol ratio of RBD-GFP following cAMP stimulation. n = 60 cells per strain from three biological replicates, with 20 cells analyzed per replicate. Data are plotted as mean ± s.e.m. **(C,D)** Representative Ras crescent assays in latrunculin-treated Ax2 and *pikF⁻* cells expressing RBD-GFP stimulated with a microcapillary containing 10 µM cAMP, indicated by the asterisk. Images from the time course are shown with corresponding kymographs below. Red arrows indicate the approximate position of the microcapillary in the kymographs. Scale bar, 5 µm. **(E,F)** Rose plots showing the angle θ between the central axis of the Ras crescent and the position of the cAMP-filled microcapillary. Bars show the proportion of measurements in each angular bin. n = 20 cells per strain across three biological replicates. **(G)** Circular variance of Ras crescent positioning in Ax2 and *pikF⁻* cells, calculated from the angle measurements shown in E and F. Data are plotted as mean ± s.d. Statistical significance was assessed by Mann–Whitney test; **P < 0.001.

### PikF is required for spatial precision of Ras activation in cAMP gradients

We then examined whether prolonged Ras activation was accompanied by altered spatial organization of Ras activity in a cAMP gradient. Latrunculin- and caffeine-treated RBD-GFP-expressing cells were stimulated with a 10 μM cAMP-filled microcapillary, and the angle between the central axis of the Ras crescent and the position of the needle was measured (Fig. 6C–G; Movies 5, 6). Caffeine treatment was used to reduce basal signaling and suppress spontaneous crescent formation. In Ax2 cells, Ras crescents formed toward the cAMP source, with crescent angles clustered around the needle axis (Fig. 6C,E; Movie 5). In *pikF⁻* cells, Ras crescents were less consistently aligned with the needle and often appeared as multiple crescent-like domains around the cortex (Fig. 6D,F; Movie 6). Circular variance analysis confirmed significantly reduced precision of Ras crescent positioning in *pikF⁻* cells compared with Ax2 (Fig. 6G). As a negative control, Ax2 cells exposed to a 0 μM cAMP-filled microcapillary did not form oriented Ras crescents under otherwise identical assay conditions (Fig. S7).

### PikG is required for endogenous cAMP oscillations and ACA loclalization

The more severe developmental phenotype of *pikG⁻* cells suggested a distinct failure in signal relay during the aggregation stage. Unlike *pikF⁻* cells, which initiated aggregation but progressed abnormally, *pikG⁻* cells failed to form streams or mounds during development (Fig. 3A; Movie 4). This defect was also evident in buffer-control needle assays, where Ax2 cells streamed and aggregated spontaneously. In contrast, *pikG⁻* cells failed to initiate coordinated streaming, aggregation, or mound formation in the absence of an external cAMP cue (Fig. S8; Movies 7, 8). We therefore examined whether *pikG⁻* cells were defective in the endogenous oscillatory signaling that drives aggregation-stage streaming.

To follow intracellular cAMP dynamics during pre-aggregation development, we used Flamindo2-mScarlet, a fluorescent cAMP biosensor whose intensity decreases as intracellular cAMP increases. Ax2, pikG⁻, and aca⁻ cells expressing Flamindo2-mScarlet were developed on non-nutrient agar and imaged during the pre-aggregation period (Fig. 7A–C; Movies 9-11). Ax2 cells formed oscillatory centers with periodic changes in Flamindo2 intensity during the hour before the onset of streaming. Autocorrelation analysis identified regular oscillatory behavior, and period measurements from the mean intensity traces showed an approximately 6 min oscillation period (Fig. 7B,C; Fig. S9A,E; Movie 9). In contrast, aca⁻ and *pikG⁻* cells did not show regular periodic changes in Flamindo2 intensity between 3 and 4 h after starvation (Fig. 7B,C; Fig. S9B,C; Movies 10, 11). Autocorrelation analysis did not identify significant oscillatory behavior in either strain, and sine-curve fitting yielded poorer fits for aca⁻ and *pikG⁻* traces than for Ax2 traces (Fig. S9A–D). Thus, *pikG⁻* cells resemble aca⁻ cells in failing to establish detectable pre-aggregation cAMP oscillations.

**Fig. 7.**
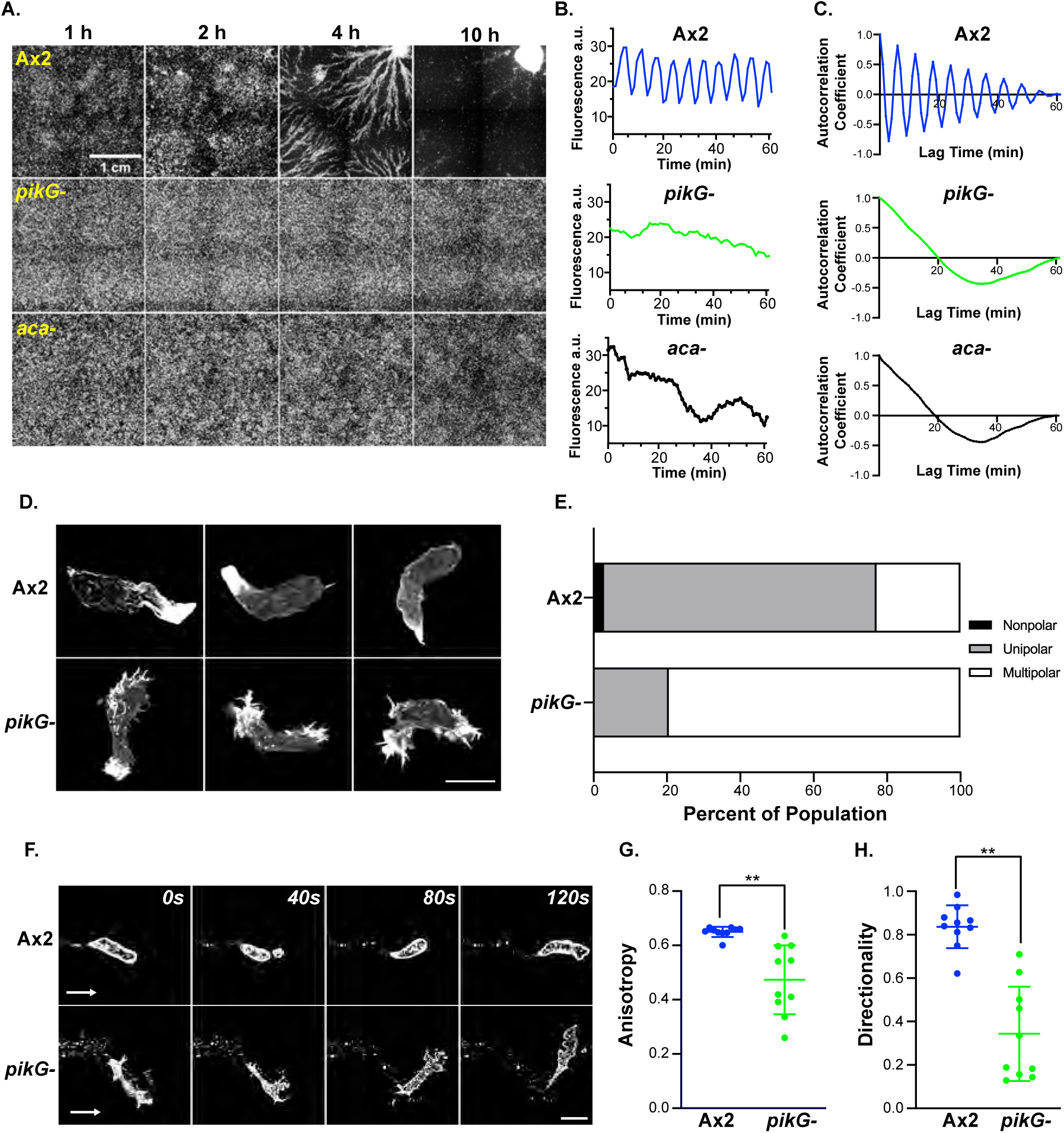
PikG is required for pre-aggregation oscillatory signaling and ACA polarity. **(A)** Representative images from developmental time courses of Ax2, *pikG⁻*, and *aca⁻* cells expressing Flamindo2-mScarlet during development on non-nutrient agar at the indicated times. **(B)** Representative fluorescence-intensity traces from oscillatory centers in Ax2, *pikG⁻*, and *aca⁻* cells during the pre-aggregation phase. **(C)** Autocorrelation analysis of the representative traces shown in B. **(D)** Representative fixed developed Ax2 and *pikG⁻* cells expressing ACA-YFP. **(E)** Quantification of ACA-YFP polarity in developed Ax2 and *pikG⁻* cells. Cells were scored as nonpolar, unipolar, or multipolar based on ACA-YFP cortical enrichment as described in Methods. n = 39 cells per strain. **(F)** Representative frames from time-lapse images of developed Ax2 and *pikG⁻* cells expressing ACA-YFP. Arrows indicate direction of cell movement. Scale bar, 10 µm. **(G)** Quantification of ACA-YFP vesicle anisotropy. n = 10 cells per strain. **(H)** Quantification of ACA-YFP vesicle directionality. n = 10 cells per strain. Statistical significance was assessed by Kruskal–Wallis test; **P < 0.001 where indicated.

ACA produces cAMP and becomes enriched at the rear of aggregation-stage cells, prompting us to examine ACA-YFP localization in developed Ax2 and *pikG*⁻ cells. In Ax2 cells, ACA localized asymmetrically, with many cells displaying unipolar enrichment (Fig. 7D,E). In *pikG⁻* cells, however, ACA-YFP was frequently enriched at multiple cortical sites rather than concentrated at a single pole, consistent with the multipolar morphology of these cells and indicating disrupted ACA polarity (Fig. 7D,E).

We next asked whether this altered ACA organization was accompanied by defects in the spatial pattern of ACA-containing vesicle secretion. Super-resolution time-lapse imaging at the coverslip surface revealed that both Ax2 and *pikG⁻* cells deposited ACA-YFP-positive vesicles, indicating that loss of PikG did not eliminate production or release of ACA-containing vesicle trails (Fig. 7F). However, the organization of these trails differed markedly between strains. Ax2 cells left relatively narrow, directionally organized ACA-YFP-positive trails, whereas *pikG⁻* cells deposited broader, less coherent trails with more frequent changes in direction (Fig. 7F; Movies 12, 13). Quantification of trail anisotropy and directionality confirmed this defect: *pikG⁻* trails had lower anisotropy and directionality scores than Ax2 trails, consistent with poorly coordinated spatial deposition of ACA-YFP-positive vesicles (Fig. 7G,H). Consistent with these measurements, representative time-lapse movies showed Ax2 cells moving in close succession along ACA-YFP-positive trails, including examples in which a following cell contacted the rear of a leader cell during streaming-like movement, whereas *pikG*⁻ cells interacted with ACA-YFP-positive trails without forming comparably coordinated streams (Movies 12, 13).Thus, PikG is not strictly required for ACA-YFP-positive vesicle deposition, but is required for the spatial organization of ACA-containing relay structures during aggregation-stage signaling.

### PikF and PikG developmental functions require conserved catalytic lysines

Finally, we tested whether the developmental defects in *pikF⁻* and *pikG⁻* cells require the predicted kinase activities of PikF and PikG. Rescue strains expressed either wild-type PikF or PikG, or kinase-dead alleles carrying mutations in the predicted catalytic lysine residues: PikF^K1119A and PikG^K1415A. Development was assayed on non-nutrient agar over 48 h using the same endpoints measured in Fig. 3, including time to mound emergence, and the timing of fruiting body formation (Fig. 8A–C).

**Fig. 8.**
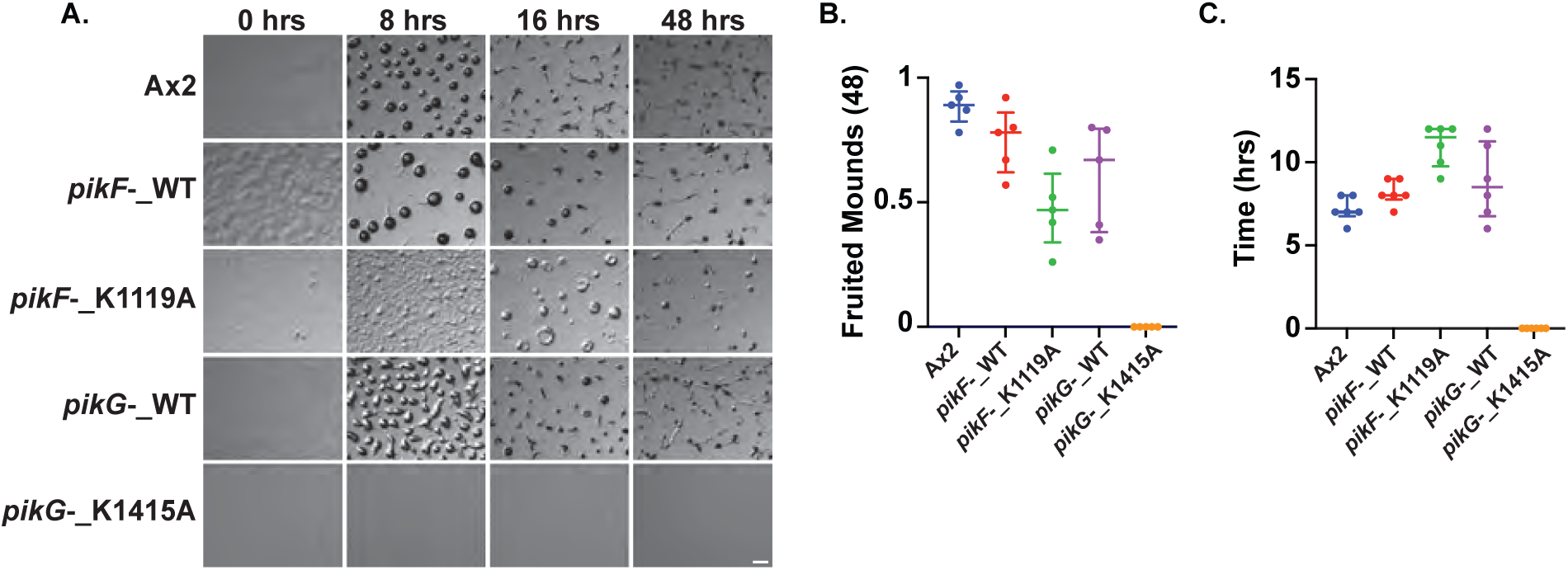
Wild-type, but not kinase-dead, PikF and PikG rescue developmental defects in *Dictyostelium*. **(A)** Representative time course of development on non-nutrient agar for the indicated strains. *pikF⁻* and *pikG⁻* cells were rescued with wild-type PikF or PikG, respectively, or with kinase-dead alleles carrying catalytic lysine mutations, PikF^K1119A and PikG^K1415A. Images were acquired over 48 h at 7.3× magnification. n = 5 independent experiments. **(B)** Fraction of mounds reaching the fruiting-body stage by 48 h. Approximately 30 mounds were scored per plate for each strain. Bars indicate mean ± s.e.m.; n = 5 biological replicates. **(C)** Time to emergence of 10% of total mounds formed by each strain during development on non-nutrient agar. Bars indicate mean ± s.e.m.; n = 5 biological replicates. Statistical significance was assessed by mixed-effects analysis. Wild-type *PikF⁻* and *PikG⁻* significantly rescued developmental progression relative to the corresponding knockout strains (P < 0.001), whereas kinase-dead PikF^K1119A and PikG^K1415A did not significantly rescue development.

Re-expression of wild-type PikF restored development in *pikF⁻* cells, producing mounds and fruiting bodies similar to Ax2 by 48 h, whereas *pikF⁻* cells expressing PikF^K1119A remained delayed and abnormal, resembling the knockout phenotype (Fig. 8A–C). Similarly, wild-type PikG restored aggregation and fruiting body formation in *pikG⁻* cells, whereas *pikG⁻* cells expressing PikG^K1415A remained aggregation-defective (Fig. 8A–C). Quantification of mound-emergence timing and the fraction of mounds reaching the fruiting-body stage confirmed that wild-type PikF and PikG restored developmental progression, whereas the corresponding kinase-dead alleles did not (Fig. 8B,C). Thus, PikF- and PikG-dependent development requires the conserved catalytic lysines necessary for PI3K activity.

## Discussion

### Atypical PI3Ks expand the PI3K signaling toolkit in *Dictyostelium*

*Dictyostelium* PI3K signaling has been studied largely through canonical Class I-like enzymes and their PI(3,4,5)P₃ output, yet several PI3K-dependent behaviors are not fully accounted for within that framework. This study identifies distinct cellular roles for three atypical PI3K-family enzymes in *Dictyostelium*. PikF constrains Ras–phosphoinositide–actin signaling during chemotaxis, PikG organizes the cAMP relay that initiates aggregation, and PikH contributes to phagocytic uptake. Together, these findings indicate that PI3K-family function is distributed across related enzymes that regulate separable outputs in protrusion, uptake, relay, and multicellular development.

Two unresolved areas in the field motivated this expanded view of *Dictyostelium* PI3K signaling. The strongest evidence that PI3K-dependent behavior extends beyond canonical Class I PI(3,4,5)P₃ production comes from studies investigating PI(3,4)P₂. PI(3,4)P₂-positive membrane domains persist after inhibitor-mediated depletion of PI(3,4,5)P₃ and remain detectable at rear membranes and macropinosome-associated compartments in PI3K1/2-null cells, suggesting that PI(3,4)P₂ organization cannot be explained solely as a PikA/PikB-dependent PI(3,4,5)P₃ turnover product (Li *et al*., 2021). A second, more indirect tension comes from studies of cAMP relay. Broad-spectrum pharmacological PI3K inhibition with LY294002 strongly impairs ACA activation and adaptation, whereas genetic loss of the major chemoattractant-responsive Class I PI3Ks PikA/PikB permits an initial ACA activation peak but disrupts adaptation (Comer and Parent, 2006). This mismatch suggests that ACA regulation may involve PI3K-dependent inputs not captured by PikA/PikB loss alone. However, this interpretation remains provisional because LY294002 can have off-target effects, including effects on TOR signaling, and because pi3kA⁻/B⁻ cells retain additional PI3K-family enzymes that could contribute to signaling. In this context, the *pikG⁻* phenotype is notable: *pikG⁻* cells fail to generate endogenous cAMP oscillations and show disrupted ACA polarity during aggregation (Fig. 7A–E; Fig. S9). Together with the PI(3,4)P₂ observations above, these findings identify PikF, PikG, and PikH as genetic entry points into PI3K-dependent behaviors not readily explained by canonical Class I PI(3,4,5)P₃ production alone.

### PikF attenuates Ras–phosphoinositide signaling and constrains protrusive activity

PikF plays a critical role in signal attenuation downstream of chemotactic stimulation. Unlike canonical Class I PI3Ks, which promote early cAMP-induced phosphoinositide signaling, PikF limits the duration and spatial spread of Ras–phosphoinositide–actin signaling after stimulation. This interpretation is supported by prolonged Ras activation, extended PH-CRAC/PIP₃ recruitment, sustained PI(3,4,5)P₃ measured by Mass ELISA, elevated cortical actin activity, and reduced directional precision during chemotaxis in *pikF⁻* cells (Fig. 2G,H; Fig. 5A–C; Fig. 6; Fig. S5D; Fig. S6). Together, these phenotypes place PikF at the level of signal attenuation rather than signal initiation.

A compelling model is that PikF helps couple PI(3,4)P₂-associated membrane dynamics to a refractory state that restrains Ras activity. This model fits the mutually inhibitory Ras–PI(3,4)P₂ polarity framework, in which Ras activity marks front-state signaling while PI(3,4)P₂ is associated with rear-state identity, protrusion suppression, and recruitment of PI(3,4)P₂-responsive RasGAPs that locally restrain Ras activity (Li *et al*., 2018). The reverse-fountain model further suggests that PI(3,4)P₂-enriched macropinosomes can give rise to vesicles that deliver PI(3,4)P₂ to the rear membrane, helping maintain a back-to-front polarity system (Li *et al*., 2021). In this context, the *pikF⁻* phenotype is consistent with a failure to properly couple PI(3,4)P₂-associated membrane dynamics to Ras attenuation: Ras activity persists, actin activity increases, and cortical signaling is no longer temporally or spatially restrained.

PikF is an attractive candidate for this role because *pikF⁻* cells show delayed PI(3,4)P₂ biosensor recovery together with prolonged Ras and PIP₃ signaling after cAMP stimulation (Fig. 5; Fig. 6A,B; Fig. S4; Fig. S5D). Importantly, *pikF⁻* cells do not simply lose acute PI(3,4,5)P₃ production; instead, they show prolonged PIP₃ signaling, placing the defect at the level of signal attenuation rather than signal initiation. This model is also supported by the structural and evolutionary features described in Figure 1: PikF retains activation-loop/basic-residue features predicted to influence phosphoinositide substrate engagement and segregates with PI3K-family enzymes related to Class II PI3Ks, which can generate PI(3,4)P₂. These data do not prove that PikF directly generates PI(3,4)P₂ or defines the relevant PI(3,4)P₂-associated refractory pool, but they strengthen the model that PikF promotes PI(3,4)P₂-linked Ras attenuation after chemotactic stimulation.

The physiological consequences of PikF loss are evident across migration, uptake, and development (Fig. 2; Fig. 3; Fig. 4). During random migration, failure to constrain Ras-driven actin activity would be expected to increase lateral protrusions and reduce persistence. During chemotaxis, the same failure would allow protrusive responses away from the cAMP source, producing the wide-angle steps and meandering paths observed in *pikF⁻* cells. The phagocytosis and macropinocytosis defects in *pikF⁻* cells (Fig. 4), suggest that this defect may extend beyond chemotactic protrusions to other actin-dependent membrane remodeling events. Macropinocytosis and large-particle uptake require spatially organized Ras–PI3K–PIP₃ signaling and coordinated actin cup closure, and PI(3,4)P₂ has been implicated at late cup stages. Thus, if PikF contributes to a PI(3,4)P₂-associated attenuation or closure pathway, impaired PikF function could reduce the efficiency of macropinocytic or phagocytic cup completion without abolishing uptake altogether. These single-cell phenotypes provide a plausible explanation for delayed mound formation and smaller aggregates, because streaming requires cells to repeatedly make directionally refined protrusive decisions in response to relay cues. The later slug and fruiting-body abnormalities may reflect cumulative effects of inefficient aggregation and disrupted tissue organization, although they may also point to additional roles for PikF-dependent phosphoinositide signaling during later morphogenesis. Thus, PikF-dependent signal attenuation appears to be required not only for accurate protrusive control in single cells, but also for the coordinated membrane and cell-movement behaviors that support multicellular development.

### PikG organizes local cAMP relay during aggregation

Whereas PikF regulates how cells attenuate and spatially constrain chemotactic signaling, PikG appears to regulate the paracrine relay system required for streaming and aggregation. *pikG⁻* cells fail to stream or aggregate and do not generate detectable Flamindo2 oscillations during pre-aggregation development (Fig. 3A; Fig. 7A–C; Fig. S9). Because Flamindo2 reports intracellular cAMP dynamics, this failure indicates that *pikG⁻* cells do not enter the periodic cAMP signaling state normally associated with relay-competent aggregation. In wild-type cells, cAMP stimulation activates a Ras–PI3K module that generates PIP₃ and recruits the PH-domain protein CRAC, promoting ACA activation, intracellular cAMP production, secretion, and relay to neighboring cells (Dormann *et al*., 2002; Comer and Parent, 2006). The absence of detectable Flamindo2 oscillations in *pikG*⁻ cells, despite their ability to respond to an external cAMP source in needle assays, therefore points to a failure in the endogenous relay circuit rather than a simple delay in morphogenesis or a general inability to sense cAMP.

The ACA-YFP and vesicle-trail data suggest that this relay defect includes a spatial-organizational component. Prior work showed that ACA becomes enriched at the rear of polarized cells and that ACA-containing vesicles can serve as relay structures capable of synthesizing and releasing cAMP (Kriebel, Barr and Parent, 2003; Kriebel *et al*., 2018). In Ax2 cells, ACA-YFP-positive trails were narrow and directionally organized, consistent with the tight head-to-tail coupling characteristic of streaming cells. In *pikG⁻* cells, ACA-YFP localization was frequently multipolar, and ACA-YFP-positive vesicles were deposited in broader, less anisotropic trails that lacked the linear persistence observed in Ax2 cells (Fig. 7D–H; Movies 12,13). Time-lapse examples further illustrate the likely consequence of this altered trail architecture: Ax2 cells often moved in close succession along ACA-YFP-positive trails left by leading cells, whereas *pikG⁻* cells produced broader, spatially disorganized ACA-YFP-positive trails and failed to maintain stable follower relationships (Movies 12,13). These observations suggest that PikG is not simply required for production or release of ACA-containing vesicles. Instead, PikG may help constrain the spatial deployment of ACA-containing relay structures, preserving the spatial coherence of paracrine relay cues during streaming. In this model, multipolar ACA organization and spatially disorganized vesicle deposition would degrade the spatial information normally provided by rear-directed relay.

How PikG constrains this spatial deployment remains unclear. One possibility is that PikG intersects with a localized PI(3,4)P₂-dependent trafficking pathway. The reverse-fountain model proposes that PI(3,4)P₂-enriched vesicles can deliver rear-state information to the back of migrating cells (Li *et al*., 2021). Although *pikG⁻* cells did not show an obvious defect in acute cAMP-stimulated PI(3,4)P₂ recovery, this does not exclude a role for PikG in spatially restricted or vesicle-associated PI(3,4)P₂ pools that help position ACA-containing relay structures.

Alternative interpretations remain possible. pikG⁻ cells could fail to amplify cAMP detection sufficiently to trigger robust relay, especially given the modest reduction in acute PH-CRAC-GFP and tPH-CynA-GFP responses (Fig. 5A–F; Fig. S4). However, their ability to migrate toward an imposed cAMP gradient with pseudopod-angle distributions similar to Ax2 argues against a gross defect in cAMP sensing (Fig. 2G,H), and the magnitude of the acute phosphoinositide defect seems unlikely, by itself, to explain the severe aggregation block. PikG could instead contribute more directly to ACA activation, adaptation, relay-circuit resetting, population-level coupling, or the biological activity of ACA-YFP-positive trails. Prior work showing that PI3K signaling can regulate both ACA activation and adaptation makes these important future directions (Comer and Parent, 2006), but the present data do not distinguish among them.

Together, the PikF and PikG phenotypes separate two kinase-dependent requirements for aggregation-stage development. PikF regulates how individual cells execute cAMP-guided movement, whereas PikG organizes the relay architecture that generates local cAMP cues for streaming. The kinase-dead rescue experiments indicate that both functions require conserved catalytic lysines, linking these distinct developmental roles to PikF and PikG kinase-domain activity (Fig. 8). This distinction explains why *pikF*⁻ cells initiate aggregation but form smaller mounds and abnormal later structures, whereas *pikG*⁻ cells fail earlier, before productive streaming and aggregation.

### PikH defines an uptake-associated branch of atypical PI3K function

PikH separates uptake-associated atypical PI3K function from developmental roles of PikF and PikG. Unlike *pikF⁻* and *pikG⁻* cells, *pikH⁻* cells resembled Ax2 in random migration, chemotaxis, development, and acute cAMP-stimulated PIP₃ and PI(3,4)P₂ dynamics (Fig. 2; Fig. 3; Fig. 5). The absence of a pronounced migration or relay phenotype is informative because it argues that the PikH branch does not contribute substantially to the acute plasma-membrane signaling events measured after cAMP stimulation. Instead, *pikH⁻* cells showed a selective defect in phagocytic uptake, while *pikG⁻* cells behaved similarly to Ax2 across the uptake assays (Fig. 4). Thus, PikH appears to define a separable uptake-associated module within the atypical PI3K family.

Wild-type PikH, but not PikH^K1586A, rescued zymosan uptake, supporting a kinase-domain-dependent role for PikH in phagocytosis (Fig. S3). Given PikH’s relationship to Vps34/Class III-like kinases, PikH may act at phagosomal, endosomal, or trafficking-associated membranes rather than in the acute plasma-membrane cAMP responses measured here. Thus, PikH defines a kinase-dependent, uptake-associated branch of atypical PI3K function.

### Future directions

An important next step is to connect each atypical PI3K to its lipid product and site of action. The kinase-dead rescue experiments show that conserved catalytic lysines are required for PikF and PikG-dependent development and PikH-dependent uptake (Fig. 8; Fig. S3A), but they do not identify the relevant phosphoinositide products. Acute recruitment of PikF, PikG, and PikH kinase domains to defined membranes, combined with live lipid biosensors, will be needed to test whether these enzymes generate distinct 3-phosphoinositide signals and where those signals act.

A broader unresolved question is how cell-autonomous polarity circuits are coupled to extracellular relay systems during collective migration. *Dictyostelium* aggregation requires cells to interpret cAMP gradients while also producing, polarizing, and releasing cAMP cues that guide their neighbors. The separation of PikF and PikG phenotypes provides a genetic entry point into this interface: PikF primarily affects the attenuation and spatial precision of intracellular protrusive signaling, whereas PikG affects the relay architecture that allows cells to generate coherent local cues. This distinction may point to a more general principle in amoeboid migration, where intracellular polarity and paracrine amplification must be coordinated to convert individual cell movement into collective guidance (Artemenko, Lampert and Devreotes, 2014; Subramanian, Majumdar and Parent, 2017).

Together, our results argue that atypical PI3K-family enzymes divide PI3K function into separable cell-biological modules. PikF constrains Ras-driven protrusive signaling, PikG organizes local cAMP relay through ACA-containing signaling structures, and PikH supports phagocytic uptake. This division of labor provides a framework for understanding how phosphoinositide kinases convert local membrane signals into coordinated changes in protrusion, uptake, relay, and multicellular development.

## Methods

### Cell lines and culture

Wild-type *Dictyostelium discoideum* Ax2 cells were obtained from the DictyBase Stock Center, with Ax2 derived from DBS0235521 (Fey et al., 2013). The *pikF⁻*, *pikG⁻*, and *pikH⁻* strains were generated in the Rob Kay laboratory by homologous recombination with insertion of a blasticidin S resistance cassette, as described previously (Hoeller and Kay, 2007). Cells were cultured axenically in HL5 medium (Formedium, UK) supplemented with vitamins and trace elements, streptomycin, and ampicillin at 21.7°C on tissue-culture-treated plastic and maintained at 40–70% confluency. All live-cell experiments were performed at 21–22°C.

Cells were transformed by electroporation using an adapted DictyBase protocol and selected in HL5 medium containing 20 µg/mL G418, 50 µg/mL hygromycin B, or both, as appropriate.

Vegetative cells expressing fluorescent constructs were incubated in KK2 buffer, containing 20 mM KH₂PO₄ and 20 mM K₂HPO₄, pH 6.4, for up to 60 min before imaging to reduce HL5-associated autofluorescence and photosensitivity.

For differentiation, cells were washed twice in development buffer, starved in suspension at 2 × 10⁷ cells/mL for 1 h, and pulsed with 100 nM cAMP every 6 min for 4–5 h. Development buffer contained 5 mM Na₂HPO₄, 5 mM KH₂PO₄, 2 mM MgSO₄, and 0.2 mM CaCl₂.

### Plasmids and rescue construct generation

pDM1203-Flamindo2-linker-mScarlet was a gift from Allyson Sgro. ACA-YFP was a gift from Carole Parent. PH-CRAC-GFP and Raf1-RBD-GFP were gifts from the Peter Devreotes laboratory. LimE-RFP was obtained from the Doug Robinson laboratory. pDM314 and pDM317 vector backbones were obtained from DictyBase.

Full-length coding sequences for *PIKF* (DDB_G0268548), *PIKG* (DDB_G0282625), and *PIKH* (DDB_G0291093) were synthesized and cloned by GenScript. Initial cloning was performed in the pDM314 backbone, and rescue constructs were subsequently generated in the GFP-tagging vector pDM317 (Veltman et al., 2009). Wild-type PikF, PikG, and PikH rescue constructs were generated alongside kinase-dead variants carrying catalytic lysine-to-alanine substitutions: PikF^K1119A, PikG^K1415A, and PikH^K1586A. All constructs were sequence verified.

### Sequence alignment and guide-tree visualization

Full-length PI3K protein sequences were obtained from DictyBase or UniProt and used for comparative sequence analysis. *Dictyostelium discoideum* sequences included the Class I PI3Ks PikA/DdPIK1 (DDB_G0278727), PikB/DdPIK2 (DDB_G0283081), and PikC/DdPIK3

(DDB_G0275011); the Class III/Vps34-like kinase PikE/DdPIK5 (DDB_G0289601); and the atypical PI3K-family proteins PikF (DDB_G0268548), PikG (DDB_G0282625), and PikH (DDB_G0291093). Human comparator sequences included Class I PI3Kα/PIK3CA (P42336), PI3Kβ/PIK3CB (B4DER4), and PI3Kγ/PIK3CG (P48736); Class II PI3K-C2α/PIK3C2A (O00443), PI3K-C2β/PIK3C2B (O00750), and PI3K-C2γ/PIK3C2G (O75747); and Class III

VPS34/PIK3C3 (Q8NEB9). Protein sequences were aligned using the Clustal Omega web server hosted by EMBL-EBI. The Clustal Omega guide tree was exported and visualized using FigTree.

### Random migration assay

Vegetative cells were harvested during exponential growth, centrifuged at 1500 rpm for 4 min, and resuspended in KK2 buffer at 1 × 10⁷ cells/mL. Cells were plated in the center of one-well chambered coverglass dishes, allowed to adhere briefly, and maintained in KK2 buffer. Random migration was recorded by phase contrast every 15 s for 1 h on a Nikon Eclipse Ti2-E microscope using a 10× 0.55 NA Plan Achromat objective. Random migration was analyzed in FIJI/ImageJ using the MTrackJ plugin. Individual cell centroids were manually tracked at 120 second intervals over the imaging period. Cell trajectories were used to calculate total path length, net displacement, migration speed, and directional persistence, defined as net displacement divided by total path length.

### Micropipette chemotaxis assay

Developed cells were plated in DB at approximately 5 × 10⁵ cells/mL in one-well coverslip chambers (Nunc Lab-Tek). Glass capillaries (BF100-58-10; Sutter Instruments) were pulled to a 0.8–1.5 µm tip and filled with 10 µM cAMP. cAMP gradients were generated by allowing a slow leak from the capillary under 10–20 hPa backpressure using a Femto microinjector. Chemotaxis toward the micropipette was imaged by phase contrast on a Nikon Eclipse Ti2-E microscope using either a 10× 0.55 NA or 40× oil 1.4 NA objective. For buffer-control assays, micropipettes containing buffer without cAMP were positioned adjacent to developed cells under otherwise identical imaging conditions.

Chemotactic alignment was quantified in FIJI/ImageJ. For each time interval, we calculated the angular deviation θ between the ideal path vector, defined as the straight-line direction toward the chemoattractant source, and the actual step vector, defined by the cell displacement between consecutive frames. Angular deviations were summarized using circular variance, calculated as 1−R, where R is the mean resultant vector length of the step-angle distribution.

Lower circular variance indicates that sequential cell steps were consistently aligned with the ideal path, whereas higher circular variance indicates more variable step directions relative to the ideal path.

### Development on non-nutrient agar

Developmental progression was assayed on 1% non-nutrient KK2 agar. For each strain, 10^7 cells were harvested during exponential growth, centrifuged at 1500 rpm for 4 min at 22°C, and washed three times with KK2 buffer. Cells were resuspended in 1 mL KK2, plated as a single central spot, and allowed to dry before imaging in a humidified chamber. Time-lapse images were acquired every 10 min for 48 h on a Leica M165 FC stereomicroscope at 7.3× magnification. Mound emergence, time to mound formation, mound width, and progression to fruiting body formation were scored from time-lapse image sequences using FIJI/ImageJ. Mounds were identified manually from time-lapse images as discrete multicellular aggregates, and approximately 30 mounds per plate were selected for width measurements or developmental-stage scoring.

### Phagocytosis Assay

For bead phagocytosis assays, 4.5 µm blue fluorescent carboxylate microspheres (Polysciences) were washed in KK2-BSA and added to cells at 50 beads per cell. Phagocytosis was stopped at the indicated time points by addition of ice-cold KK2-BSA, and cells were washed to remove uningested beads.

For zymosan phagocytosis assays, cells were harvested during exponential growth, washed twice in KK2 buffer, resuspended in KK2, and shaken for 60 min before addition of TRITC-labeled zymosan particles (Thermo Fisher Scientific). Phagocytosis was stopped at the indicated time points by addition of ice-cold KK2, and cells were washed to remove uningested particles before imaging.

For both assays, cells were imaged by brightfield and epifluorescence microscopy on a Nikon Eclipse Ti2-E microscope using a 40× 1.4 NA objective. Internalized particles were identified from paired brightfield and fluorescence images.

### Macropinocytosis assay

Cells were washed twice in KK2 buffer and starved for 30 min with shaking. Cells were plated in Nunc Lab-Tek II eight-well chambered coverglass slides and allowed to adhere briefly before addition of Alexa Fluor 647–dextran. Dextran uptake was imaged every 1 min for 1 h on a Zeiss LSM 980 laser scanning confocal microscope with Airyscan 2 using a 40× 1.4 NA objective Macropinocytosis was quantified in FIJI/ImageJ by measuring intracellular Alexa Fluor 647–dextran fluorescence intensity over time. Mean fluorescence intensity was plotted over time, and statistical analysis was performed in GraphPad Prism.

### Uniform cAMP stimulation assay

Cells were developed as described above, plated at 5 × 10⁴–1 × 10⁵ cells per well in eight-well chambered coverglass slides (Nunc Lab-Tek), and allowed to adhere for 10 min before imaging. To minimize stimulus-induced shape changes, cells were treated with 5 µM latrunculin A for 5 min. Cells were imaged on a Zeiss LSM 980 laser scanning confocal microscope with Airyscan 2 using a 40× 1.4 NA objective and stimulated with 10 µM cAMP during image acquisition. Biosensor redistribution was imaged every 2 s before and after stimulation. This assay was used to quantify cAMP-stimulated dynamics of PH-CRAC-GFP, tPH-CynA-GFP, and Raf1-RBD-GFP.

Biosensor redistribution was quantified in FIJI/ImageJ as the membrane-to-cytosol fluorescence ratio. Membrane and cytosolic regions were defined from binary masks, background-subtracted fluorescence was measured in each region, and membrane intensity was divided by cytosolic intensity at each time point. Ratios were normalized to the prestimulation baseline, defined as the average of five prestimulation frames, where indicated. Peak response was defined as the maximum absolute deviation from baseline after cAMP addition. For PH-CRAC-GFP, recovery time was defined as the first post-peak time point at which the ratio returned to baseline. For tPH-CynA-GFP, time to 50% recovery was defined as the first post-peak time point at which the signal recovered halfway from peak response toward baseline. Recovery times were reported relative to cAMP stimulation rather than the start of image acquisition. Recovery distributions were analyzed by Kaplan–Meier survival analysis with log-rank comparisons. Peak amplitudes were compared by Kruskal–Wallis test with Dunn’s multiple-comparisons correction.

### Lipid extraction and PIP Mass ELISA

Developed cells were collected, resuspended in DB at 5 × 10⁷ cells/mL, and stimulated with 10 µM cAMP for the indicated times or treated with DB for unstimulated controls. Stimulations were stopped by addition of ice-cold 0.7 M trichloroacetic acid.

Lipids were isolated by sequential acidic extraction. Cell pellets were washed in 5% trichloroacetic acid containing 1 mM EDTA, neutral lipids were removed with methanol:chloroform, and acidic lipids were extracted with methanol:chloroform:HCl. Organic phases were collected, dried by SpeedVac, and stored at −20°C until analysis.

PI(3,4,5)P₃, PI(3,4)P₂, and PI3P levels were measured using Echelon PIP Mass ELISA kits according to the manufacturer’s instructions. Dried lipid extracts were resuspended in the supplied assay buffer, sonicated, and assayed in duplicate alongside purified PIP standards. Absorbance was measured at 450 nm using a SpectraMax plate reader, and lipid abundance was calculated from standard curves. Lipid abundance was normalized to cell number for each extraction. For time-course stimulation experiments, fold change at each time point was calculated relative to the unstimulated 0 s value from the same independent extraction.

### Peripheral actin imaging

Vegetative Ax2 and *pikF⁻* cells expressing LimE-RFP were plated in chambered coverglass dishes and imaged for 500 s on a Zeiss LSM 980 laser scanning confocal microscope with Airyscan 2 using a 40× 1.4 NA objective to monitor cortical actin dynamics. Peripheral LimE-RFP patches were quantified from time-lapse images in FIJI/ImageJ. Cortical fluorescence enrichments were scored over time, and the number of peripheral actin patches per cell was recorded.

### Ras crescent assay

Raf1-RBD-GFP-expressing cells were developed as described above. Cells were resuspended in KK2, plated in chambered coverglass dishes, and treated with 5 µM latrunculin A and 1 mM caffeine before stimulation. Cells were stimulated with a micropipette containing either 10 µM cAMP or 0 µM cAMP control positioned adjacent to the cells. RBD-GFP dynamics were imaged every 2 s on a Nikon Eclipse Ti2-E microscope equipped with a C2 confocal module using a 63× 1.4 NA objective.

Ras crescent orientation was measured in FIJI/ImageJ to quantify how closely Ras activation was oriented toward the cAMP source. For each frame, a reference line representing the most direct path to the needle was drawn from the cell centroid to the cAMP-filled micropipette tip. Each discrete Raf1-RBD-GFP-enriched cortical crescent was identified, and its position was defined by the crescent centroid. The angle ϕ between each crescent centroid and the reference line to the needle was measured. When multiple crescents were present in a single cell, each crescent was measured separately and included as a crescent-level angular measurement.

### Flamindo2 oscillatory response assay

Ax2, *pikG⁻*, and *aca*⁻ cells expressing Flamindo2-mScarlet were harvested during exponential growth, washed twice in KK2 buffer, and plated on ultrathin non-nutrient agar at 5 × 10⁶ cells/mL. Cells were developed under humidified, sealed imaging conditions and imaged on a Nikon Eclipse Ti2-E microscope at 100× magnification. Images were acquired every 60 s for 8 h.

For Ax2 cells, fields containing aggregation centers were analyzed beginning 60 min before visible streaming. Because *pikG⁻* and *aca*⁻ cells did not form aggregation centers, fields were selected randomly by an observer blinded to experimental condition, and analysis began 4 h after starvation.

Mean Flamindo2-mScarlet fluorescence intensity was measured over time in FIJI/ImageJ using the Time Series Analyzer plugin. Intensity traces were fit in GraphPad Prism using an oscillatory function to estimate amplitude and period, with goodness of fit assessed by R². Oscillation periods were also estimated using a custom Python-based autocorrelation analysis.

Fluorescence traces were mean-centered, autocorrelated, and normalized to the zero-lag value. The oscillation period was defined as the time lag of the first non-zero autocorrelation peak.

Traces without a detectable autocorrelation peak were classified as non-oscillatory.

### ACA-YFP polarity assay

Developed Ax2 and *pikG⁻* cells expressing ACA-YFP were plated on 13 mm borosilicate coverslips and fixed in 4% paraformaldehyde/PBS. Cells were imaged on a Zeiss LSM 980 laser scanning confocal microscope with Airyscan 2 using a 60× 1.4 NA objective. ACA-YFP polarity was quantified from fluorescence intensity profiles along the cell perimeter. An ACA-YFP pole was defined operationally as a cortical enrichment that spanned at least 10% of the total cell perimeter, was separated from neighboring poles by at least 10% of the perimeter, remained at least threefold above the minimum perimeter intensity, and reached at least 50% of the maximum perimeter intensity.

### ACA-YFP vesicle trail directionality and anisotropy assay

Developed Ax2 and *pikG⁻* cells expressing ACA-YFP were plated at 5 × 10⁴ cells/well in 500 µL DB in Nunc Lab-Tek II chambered coverglass slides. Cells were imaged every 10 s for 10 min on a Zeiss LSM 980 laser scanning confocal microscope with Airyscan 2 using a 40× 1.4 NA objective. Vesicle x-y coordinates were recorded in order of secretion. Vesicle trail directionality is a persistence measure calculated as net displacement divided by total path length. Vesicle trail anisotropy was calculated from the variance in vesicle position along the x- and y-axes and the x-y covariance. Eigenvalues were calculated as:

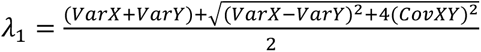 and 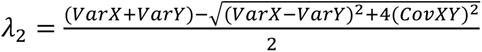 are used in the anisotropy equation 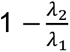 where values closest to 1 represent the most anisotropic trails and values closest to 0 represent the most isotropic trails.

### Membrane kymograph generation

Where membrane kymographs are shown, cell masks were generated from fluorescent images in FIJI/ImageJ and used to define cell boundaries for kymograph generation with Membrane Kymograph Generator (Banerjee et al., 2026).

### Statistical analysis

Statistical analyses were performed in GraphPad Prism. Statistical tests were selected based on the experimental design and are indicated in the corresponding figure legends. For comparisons among multiple strains, Kruskal–Wallis tests with Dunn’s multiple-comparisons correction or mixed-effects models were used as indicated. Circular data were analyzed using circular variance. *P*-values less than 0.05 were considered statistically significant. Sample sizes, replicate numbers, definitions of *n*, and error bars are reported in the figure legends.

## Acknowledgements

We thank A.K. Mohammadi for critical reading of the manuscript and C. Dovey for helpful discussions. The authors declare no competing or financial interests. This work was supported by the National Institutes of Health National Institute of General Medical Sciences grant 1R15GM143733-01 awarded to M.E. and by a National Science Foundation Major Research Instrumentation award DBI-2117798 awarded to S. Kim.

---

**Fig. S1.**
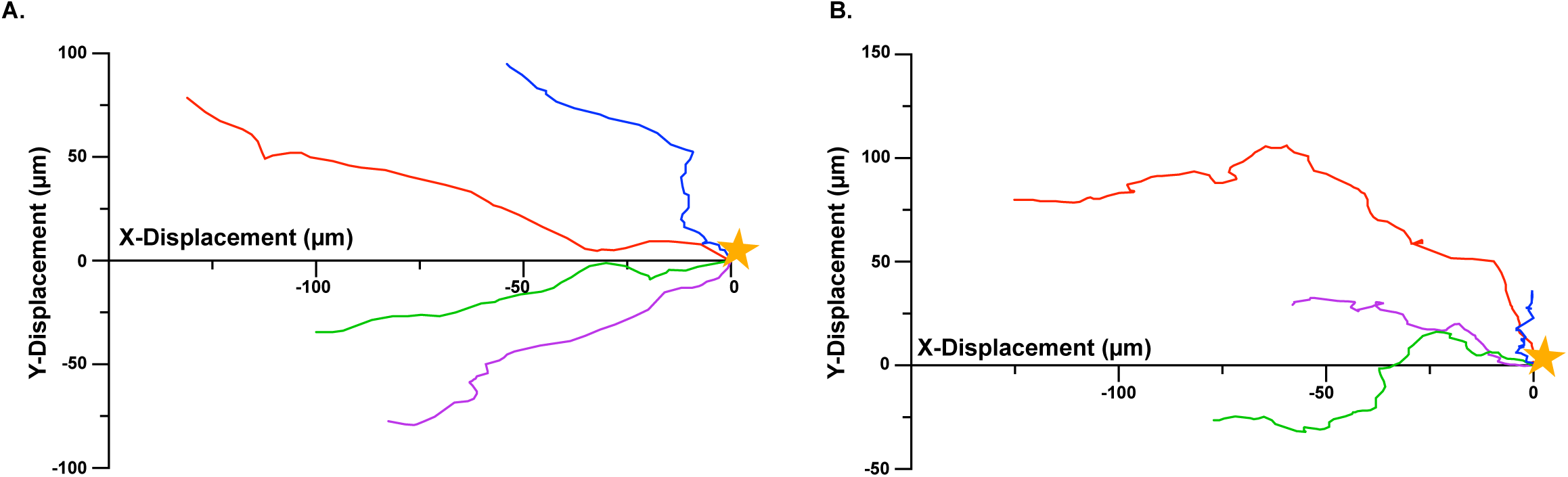
Representative directed migration trajectories of Ax2 and *pikF⁻* cells. **(A)** Four representative Ax2 cell trajectories from the directed cell migration assay. **(B)** Four representative *pikF⁻* cell trajectories from the same assay. These trajectories are from the same cells analyzed in Fig. 2G. The gold star indicates the position of the microcapillary needle.

**Fig. S2.**
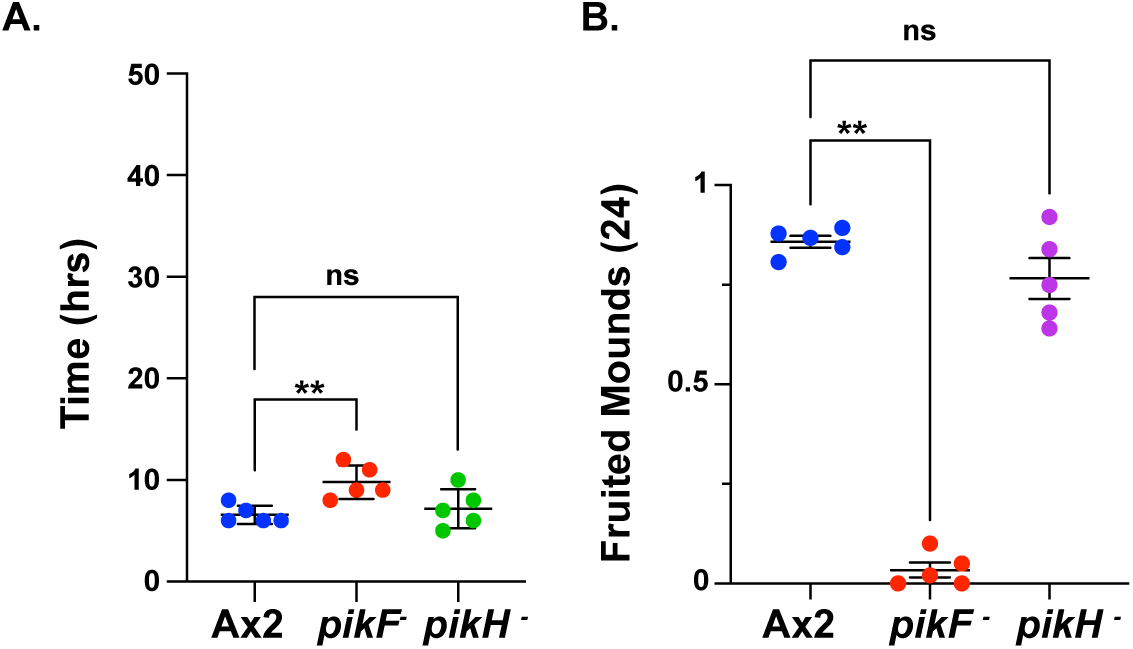
Quantification of developmental timing in Ax2 and atypical PI3K mutants. **(A)** Time to emergence of 10% of total mounds formed by Ax2, *pikF⁻*, *pikG⁻*, and *pikH⁻* cells developing on non-nutrient agar. Five biological replicates were analyzed. **(B)** Fraction of mounds reaching the fruiting-body stage by 24 h. Five biological replicates were analyzed, with approximately 30 mounds scored per plate. Bars indicate mean ± s.e.m. Statistical significance was assessed by Kruskal-Wallis test; *P < 0.001 where indicated.

**Fig. S3.**
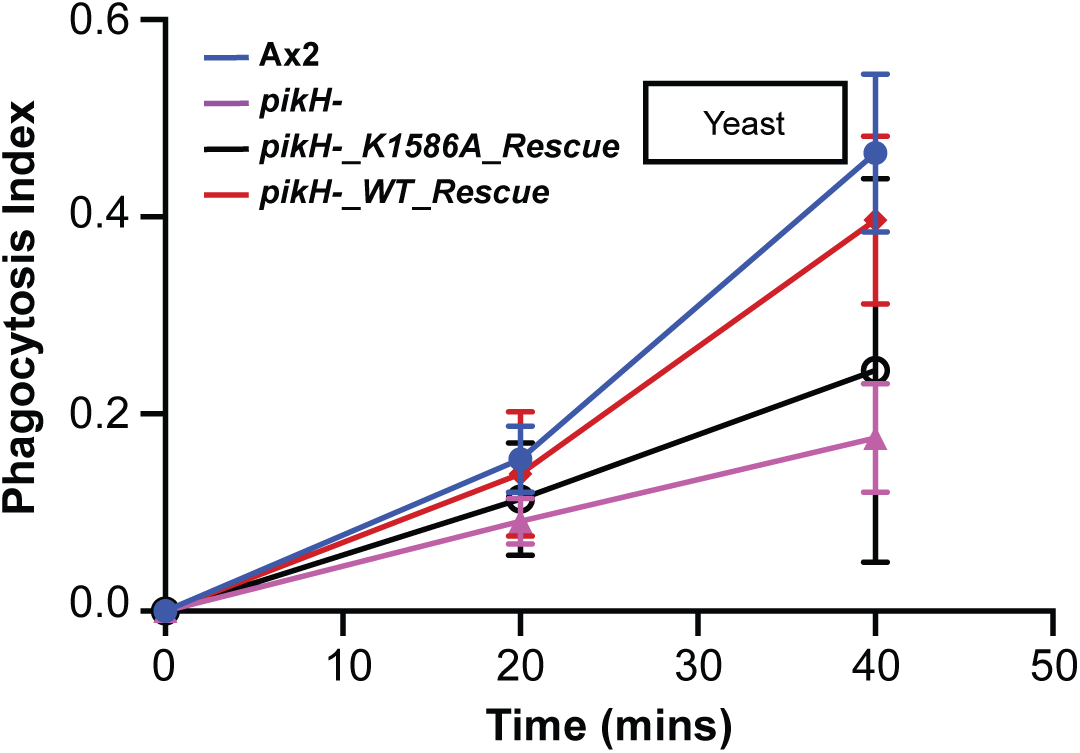
Wild-type, but not kinase-dead, PikH rescues the phagocytic uptake defect in pikH⁻ cells. Phagocytosis index for uptake of fluorescent *Saccharomyces cerevisiae* particles by the indicated strains. Re-expression of wild-type PikH restores uptake in *pikH⁻* cells, whereas the kinase-dead PikH^K1586A allele fails to rescue the uptake defect. Bars indicate mean ± s.e.m. from two biological replicates.

**Fig. S4.**
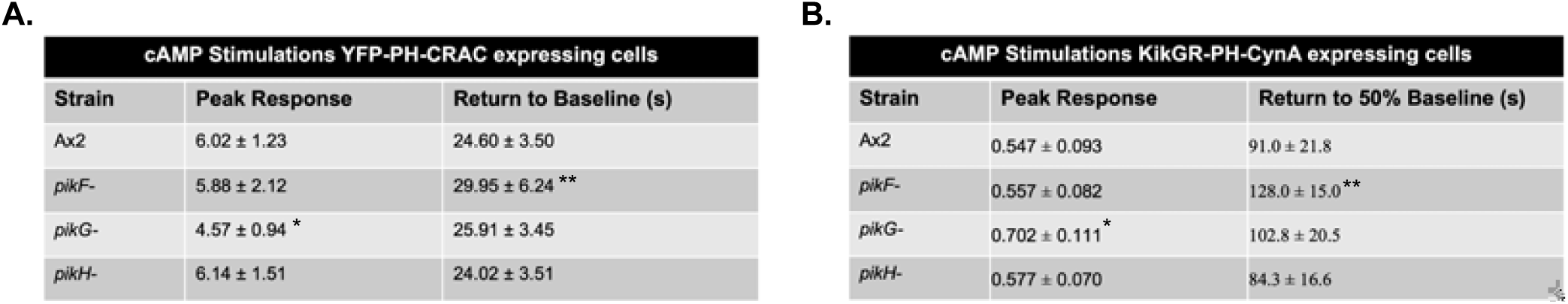
Quantification of PH-CRAC-GFP and tPH-CynA-GFP responses shown in Fig. 5. **(A)** Quantification of PH-CRAC-GFP peak response and time to return to prestimulation baseline in Ax2, *pikF⁻*, *pikG⁻*, and *pikH⁻* cells following uniform cAMP stimulation, based on the data shown in Fig. 5C. **(B)** Quantification of tPH-CynA-GFP peak response and time to 50% recovery in Ax2, *pikF⁻*, *pikG⁻*, and *pikH⁻* cells following uniform cAMP stimulation, based on the data shown in Fig. 5F. n = 60 cells per strain from three biological replicates, with 20 cells analyzed per replicate. Recovery times in S4 are reported as time after cAMP stimulation, whereas traces in Fig. 5C,F are plotted on the full acquisition timeline with cAMP added at 10 s. Peak amplitudes were compared by Kruskal–Wallis test with Dunn’s post hoc correction. PH-CRAC-GFP recovery was analyzed as time to return to prestimulation baseline. tPH-CynA-GFP recovery was analyzed as time to 50% recovery, defined as the first post-peak time point at which the signal had recovered halfway from its peak response toward the prestimulation baseline. Recovery-time distributions were analyzed using Kaplan–Meier survival analysis with log-rank comparison. *P < 0.05 where indicated.

**Fig. S5.**
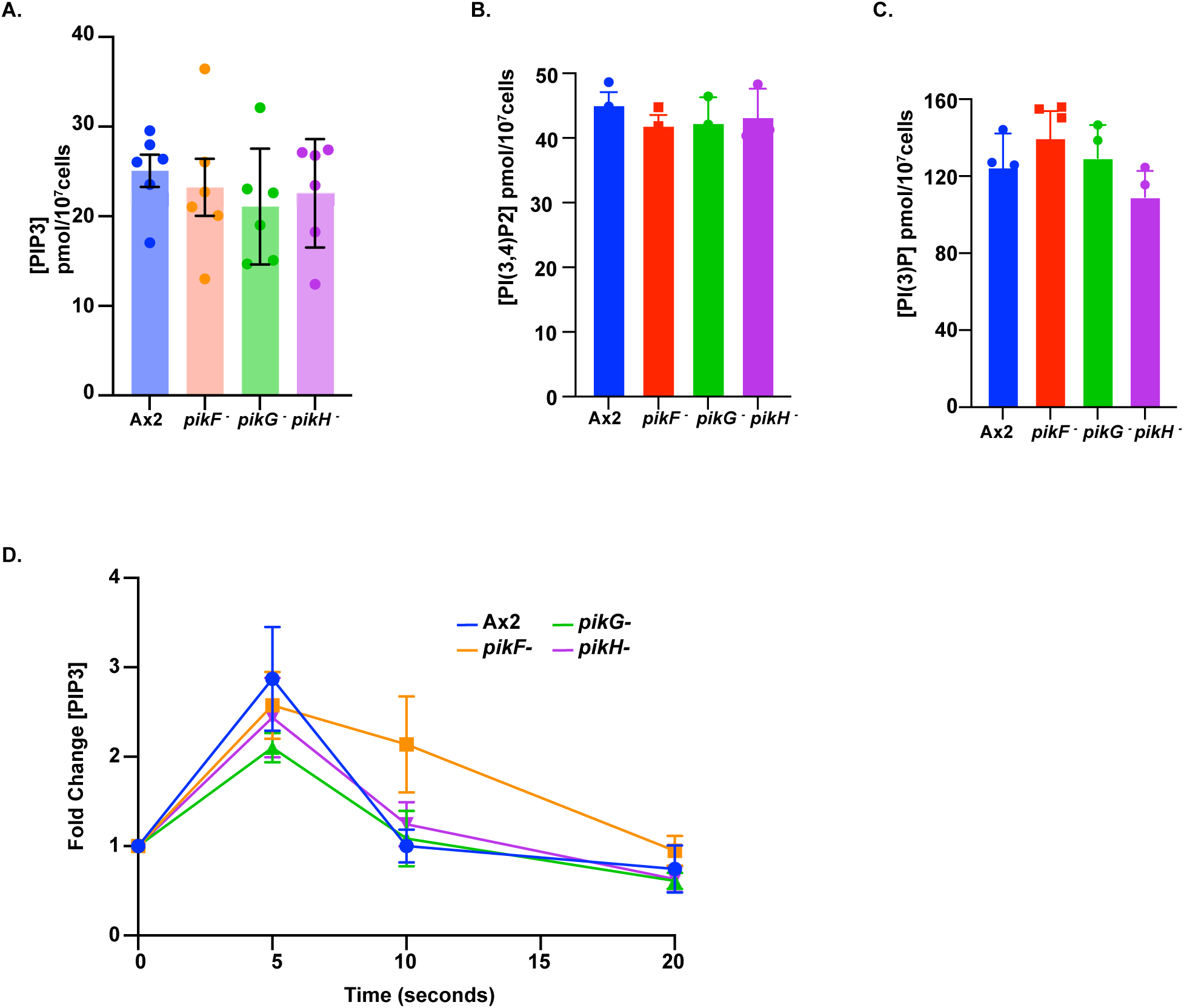
Basal phosphoinositide levels and cAMP-stimulated PI(3,4,5)P₃ dynamics in Ax2 and PI3K mutant cells. **(A)** Basal PI(3,4,5)P₃ levels in developed Ax2, *pikF⁻*, *pikG⁻*, and *pikH⁻* cells, normalized to cell number and quantified by MASS ELISA. Mean ± s.e.m.; n = 6 independent lipid extractions. **(B)** Basal PI(3,4)P₂ levels in developed Ax2, *pikF⁻*, *pikG⁻*, and *pikH⁻* cells, normalized to cell number and quantified by MASS ELISA. Mean ± s.e.m.; n = 4 independent lipid extractions. **(C)** Basal PI3P levels in developed Ax2, *pikF⁻*, *pikG⁻*, and *pikH⁻* cells, normalized to cell number and quantified by MASS ELISA. Mean ± s.e.m.; n = 3 independent lipid extractions. For A–C, basal lipid levels were not significantly different across strains by Kruskal–Wallis test with Dunn’s post hoc correction. **(D)** Relative change in PI(3,4,5)P₃ following cAMP stimulation in developed Ax2, *pikF⁻*, *pikG⁻*, and *pikH⁻* cells, quantified by MASS ELISA. Fold change was calculated relative to the unstimulated value for each strain within each independent extraction. Mean ± s.e.m.; n = 3 independent lipid extractions. Time-course data were analyzed by mixed-effects model; *pikF⁻* differed significantly from Ax2 over time, P < 0.03.

**Fig. S6.**
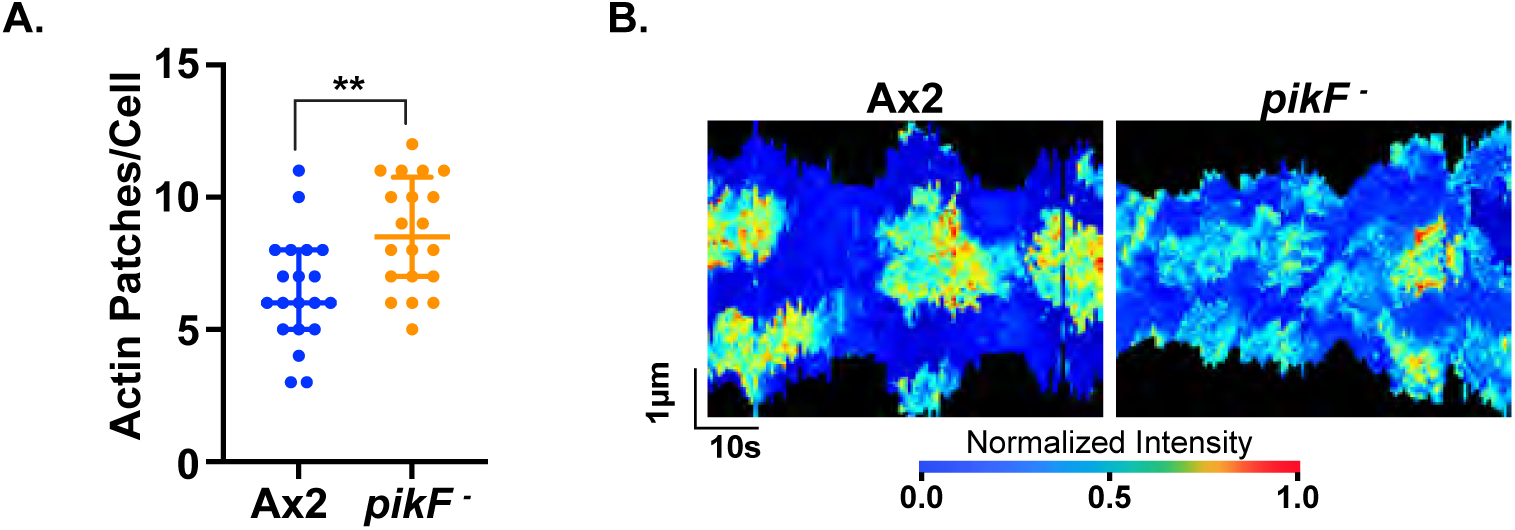
*pikF⁻* cells show increased peripheral actin activity. **(A)** Quantification of peripheral actin patches in Ax2 and *pikF⁻* cells expressing LimE-RFP. Cells were imaged for 500 s, and peripheral actin patches were scored as described in Methods. n = 20 cells per strain across three biological replicates. **(B)** Representative kymographs of cells quantified in A showing the distribution of peripheral actin patches over time. Statistical significance was assessed by Mann-Whitney test; **P < 0.001.

**Fig. S7.**
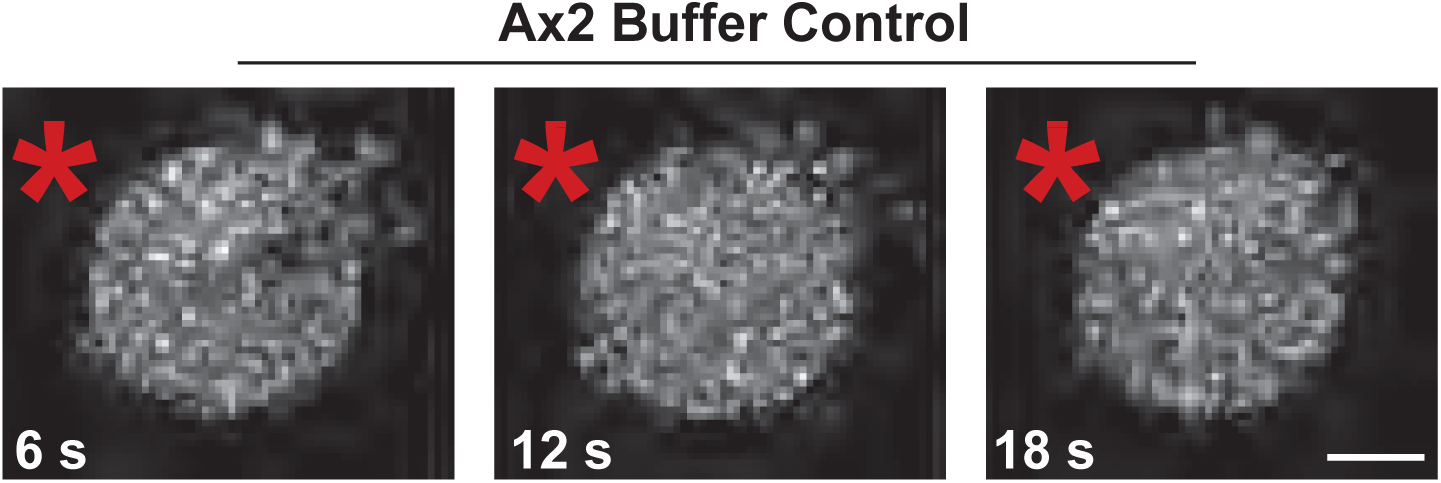
Negative control for Ras crescent assay. **(A,B)** Representative Ras crescent assays in latrunculin-treated AX2 cells expressing RBD-GFP and exposed to a 0 μM cAMP-filled microcapillary (asterisk). Red asterisk indicates the approximate needle position. Conditions are otherwise as in Fig. 5C,D. Scale bar: 5 μm.

**Fig S8.**
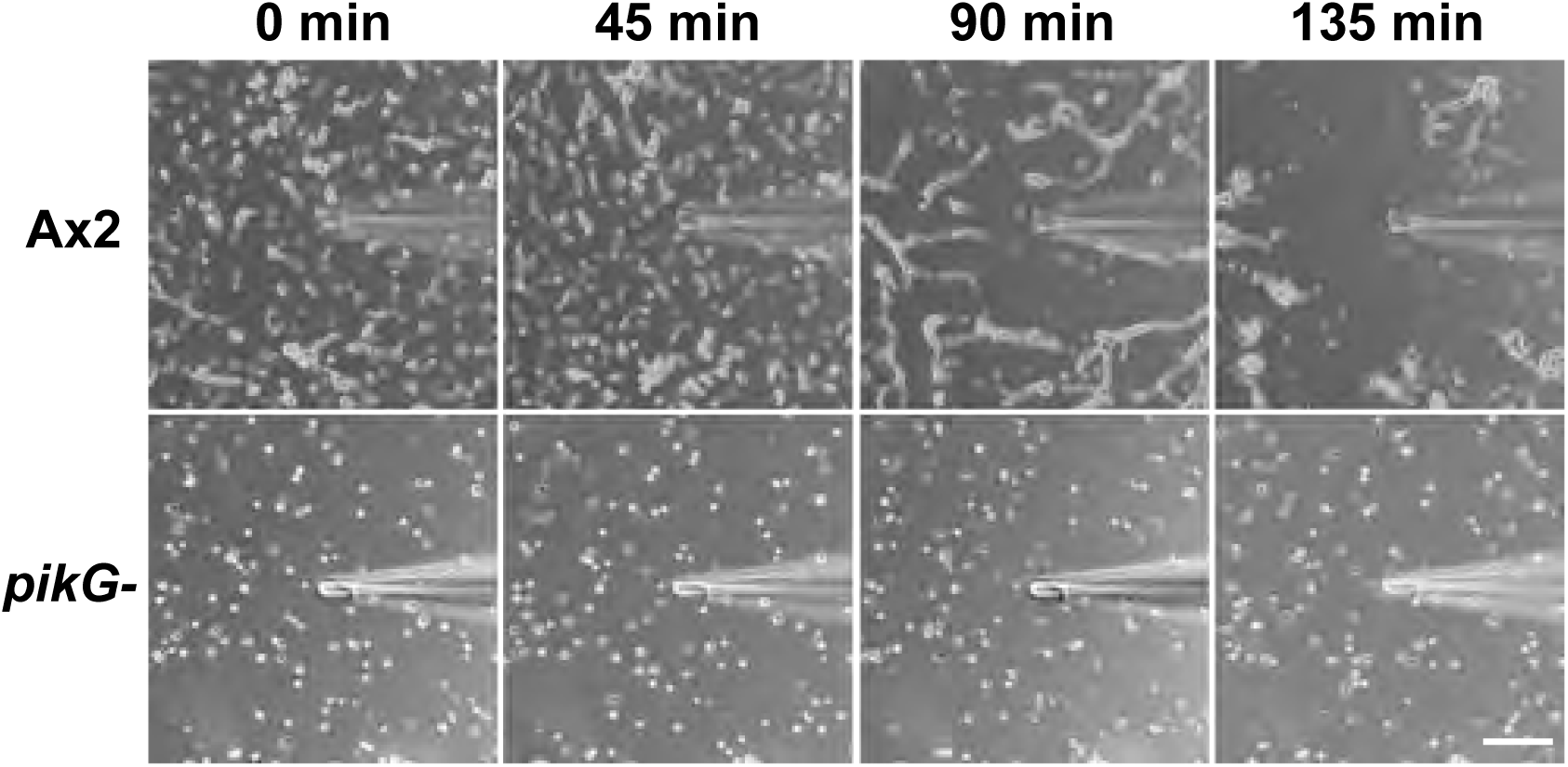
*pikG⁻* cells fail to stream in the absence of an external cAMP stimulus. Time-lapse montages of AX2 and *pikG⁻* cells developed under identical conditions and exposed to a buffer-containing microcapillary as a control for the directed migration assay. AX2 cells exhibited spontaneous streaming and aggregation, whereas *pikG⁻* cells failed to initiate coordinated streaming, aggregation or mound formation. Representative of n=3 independent experiments. Scale bar: 100 μm.

**Fig. S9.**
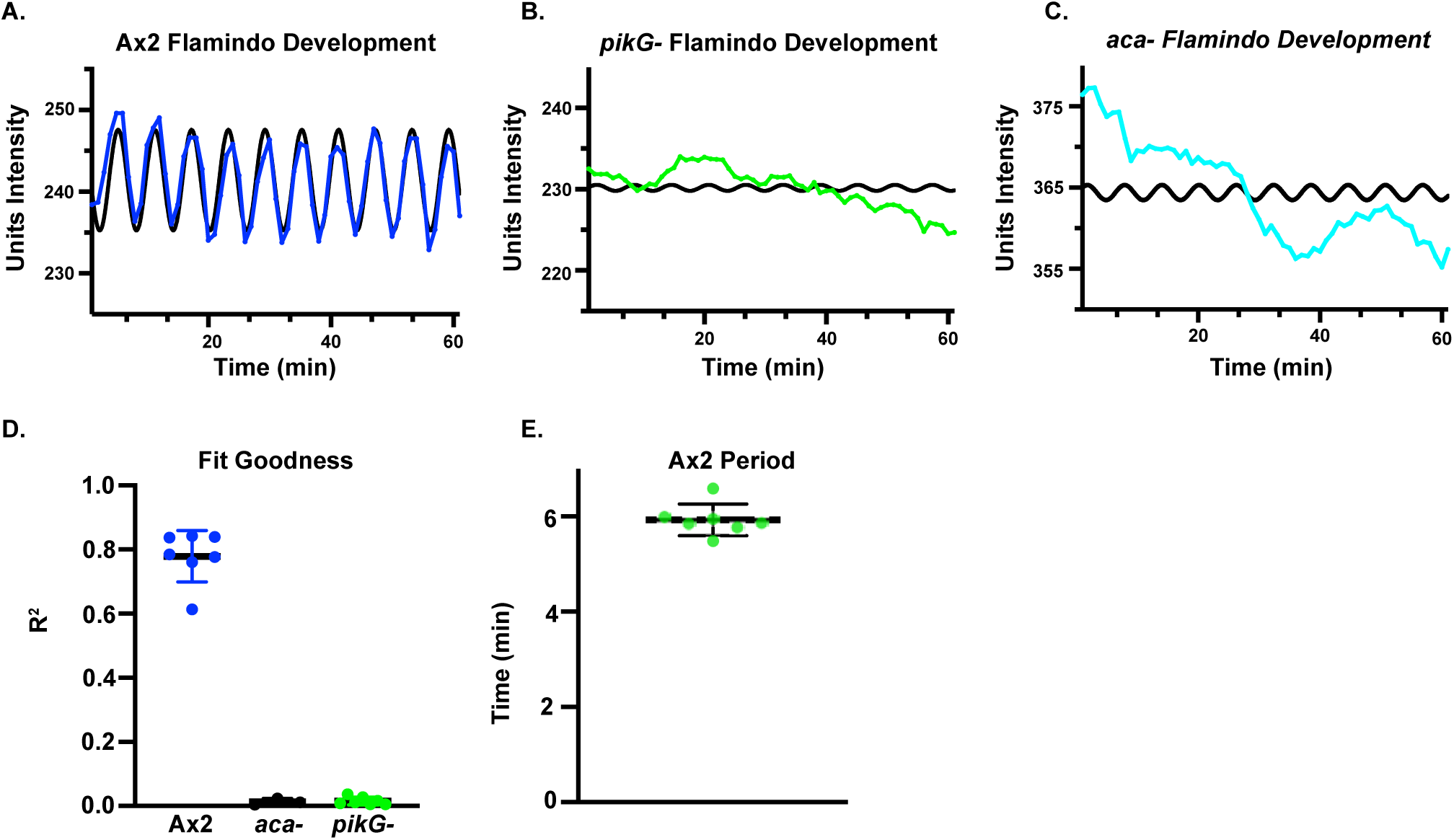
Sine curve fitting analysis of Flamindo2 oscillations. **(A–C)** Mean intensity traces from representative oscillatory centers in AX2 **(A)**, aca⁻ **(B)** and *pikG⁻* **(C)** cells, shown with fitted sine curves (black). AX2 traces are from the hour before streaming; aca⁻ and *pikG⁻* traces are from 3–4 h after starvation. **(D)** Goodness-of-fit (R²) values for sine curve fits across AX2 (n=7), aca⁻ (n=3) and *pikG⁻* (n=7) replicates. Error bars indicate median ± s.d. **(E)** Scatter plot of oscillation periods in AX2 (n=7), calculated from peaks in the mean intensity traces. Error bars indicate median ± s.d.

**Movie 1. Micropipette chemotaxis assay of developed AX2 and *pikF⁻* cells.**

Time-lapse movie of developed cells migrating toward a micropipette containing 10 µM cAMP. AX2 cells are shown on the left and *pikF⁻* cells on the right. Images were acquired on a Nikon Eclipse Ti2-E microscope using a 40×/1.4 NA oil-immersion objective at 6 frames per second and are played back at 25 frames per second. Scale bar, 4 µm.

**Movie 2. Micropipette chemotaxis assay of developed AX2 cells.**

Time-lapse movie of developed AX2 cells migrating toward a micropipette containing 10 µM cAMP. Images were acquired on a Nikon Eclipse Ti2-E microscope using a 40×/1.4 NA oil-immersion objective at 6 frames per second and are played back at 10 frames per second. Scale bar, 4 µm.

**Movie 3. Micropipette chemotaxis assay of developed *pikG⁻* cells.**

Time-lapse movie of developed *pikG⁻* cells migrating toward a micropipette containing 10 µM cAMP. Images were acquired on a Nikon Eclipse Ti2-E microscope using a 40×/1.4 NA oil-immersion objective at 6 frames per second and are played back at 10 frames per second. Scale bar, 4 µm.

**Movie 4. Development on non-nutrient agar.**

Time-lapse movie showing development of the indicated *Dictyostelium* strains on 1% non-nutrient KK2 agar. Strains are labeled in the movie. Images were acquired every 10 min for 48 h at 7.3x magnification and are played back at 15 frames per second. Scale bar, 500 µm.

**Movie 5. Ras crescent assay in developed AX2 cells expressing Raf1-RBD-GFP.**

Time-lapse movie of developed AX2 cells expressing Raf1-RBD-GFP in response to a micropipette containing 10 µM cAMP. The position of the needle is indicated by an asterisk. Images were acquired every 6 s on a Nikon Eclipse Ti2-E microscope equipped with a C2 confocal module using a 63×/1.4 NA objective and are played back at 10 frames per second. Scale bar, 2 µm.

**Movie 6. Ras crescent assay in developed *pikF⁻* cells expressing Raf1-RBD-GFP.**

Time-lapse movie of developed *pikF⁻* cells expressing Raf1-RBD-GFP in response to a micropipette containing 10 µM cAMP. The position of the needle is indicated by an asterisk. Images were acquired every 6 s on a Nikon Eclipse Ti2-E microscope equipped with a C2 confocal module using a 63×/1.4 NA objective and are played back at 10 frames per second. Scale bar, 2 µm.

**Movie 7. Buffer-control micropipette assay of developed AX2 cells.**

Time-lapse movie of developed AX2 cells exposed to a micropipette containing 0 µM cAMP. Images were acquired on a Nikon Eclipse Ti2-E microscope using a 10×/0.55 NA objective at 6 frames per second and are played back at 10 frames per second.

**Movie 8. Buffer-control micropipette assay of developed *pikG⁻* cells.**

Time-lapse movie of developed *pikG⁻* cells exposed to a micropipette containing 0 µM cAMP. Images were acquired on a Nikon Eclipse Ti2-E microscope using a 10×/0.55 NA objective at 6 frames per second and are played back at 10 frames per second.

**Movie 9. Time-lapse imaging of developed AX2 cells expressing Flamindo2-mScarlet.** Time-lapse movie of developed AX2 cells expressing Flamindo2-mScarlet. Images were acquired every 60 s for 8 h on a Nikon Eclipse Ti2-E microscope at 5× magnification. Scale bar, 500 µm.

**Movie 10. Time-lapse imaging of developed *pikG⁻* cells expressing Flamindo2-mScarlet.**

Time-lapse movie of developed *pikG⁻* cells expressing Flamindo2-mScarlet. Images were acquired every 60 s for 8 h on a Nikon Eclipse Ti2-E microscope at 5× magnification. Scale bar, 500 µm.

**Movie 11. Time-lapse imaging of developed aca⁻ cells expressing Flamindo2-mScarlet.**

Time-lapse movie of developed aca⁻ cells expressing Flamindo2-mScarlet. Images were acquired every 60 s for 8 h on a Nikon Eclipse Ti2-E microscope at 5× magnification. Scale bar, 500 µm.

**Movie 12. Streaming behavior of developed AX2 cells expressing ACA-YFP.**

Time-lapse movie of developed AX2 cells expressing ACA-YFP. Images were acquired every 10 s on a Zeiss LSM 980 with Airyscan using a 63×/1.40 NA oil-immersion objective and are played back at 5 frames per second. Scale bar, 5 µm.

**Movie 13. Streaming behavior of developed *pikG⁻* cells expressing ACA-YFP.**

Time-lapse movie of developed *pikG⁻* cells expressing ACA-YFP. Images were acquired every 10 s on a Zeiss LSM 980 with Airyscan using a 63×/1.40 NA oil-immersion objective and are played back at 5 frames per second. Scale bar, 5 µm.

## References

1. Andrew, N. and Insall, R.H. (2007) “Chemotaxis in shallow gradients is mediated independently of PtdIns 3-kinase by biased choices between random protrusions,” Nature Cell Biology, 9(2), pp. 193–200. Available at: 10.1038/ncb1536.

2. Artemenko, Y., Lampert, T.J. and Devreotes, P.N. (2014) “Moving towards a paradigm: common mechanisms of chemotactic signaling in Dictyostelium and mammalian leukocytes,” Cellular and Molecular Life Sciences, 71(19), pp. 3711–3747. Available at: 10.1007/s00018-014-1638-8.

3. Balla, T. (2013) “Phosphoinositides: Tiny lipids with giant impact on cell regulation,” Physiological Reviews, 93, pp. 1019–1137.

4. Brown, J.R. and Auger, K.R. (2011) “Phylogenomics of phosphoinositide lipid kinases: perspectives on the evolution of second messenger signaling and drug discovery,” BMC Evolutionary Biology, 11(1), p. 4. Available at: 10.1186/1471-2148-11-4.

5. Charest, P.G. and Firtel, R.A. (2006) “Feedback signaling controls leading-edge formation during chemotaxis,” Current Opinion in Genetics & Development, 16(4), pp. 339–347. Available at: 10.1016/j.gde.2006.06.016.

6. Comer, F.I. and Parent, C.A. (2006) “Phosphoinositide 3-Kinase Activity Controls the Chemoattractant-mediated Activation and Adaptation of Adenylyl Cyclase,” Molecular Biology of the Cell, 17(1), pp. 357–366. Available at: 10.1091/mbc.e05-08-0781.

7. Devreotes, P.N. et al. (2017) “Excitable Signal Transduction Networks in Directed Cell Migration.,” Annual review of cell and developmental biology, 33, pp. 103–125. Available at: 10.1146/annurev-cellbio-100616-060739.

8. Dormann, D. et al. (2002) “Visualizing PI3 kinase-mediated cell-cell signaling during Dictyostelium development.,” Current biology : CB, 12(14), pp. 1178–1188. Available at: 10.1016/s0960-9822(02)00950-8.

9. Dormann, D. and Weijer, C.J. (2003) “Chemotactic cell movement during development,” Current Opinion in Genetics & Development, 13(4), pp. 358–364. Available at: 10.1016/s0959-437x(03)00087-x.

10. Engelman, J.A., Luo, J. and Cantley, L.C. (2006) “The evolution of phosphatidylinositol 3-kinases as regulators of growth and metabolism,” Nature Reviews Genetics, 7(8), pp. 606–619. Available at: 10.1038/nrg1879.

11. Ferguson, G.J. et al. (2007) “PI(3)Kγ has an important context-dependent role in neutrophil chemokinesis,” Nature Cell Biology, 9(1), pp. 86–91. Available at: 10.1038/ncb1517.

12. Friedl, P. and Wolf, K. (2010) “Plasticity of cell migration: a multiscale tuning model,” The Journal of Cell Biology, 188(1), pp. 11–19. Available at: 10.1083/jcb.200909003.

13. Funamoto, S. et al. (2002) “Spatial and Temporal Regulation of 3-Phosphoinositides by PI 3-Kinase and PTEN Mediates Chemotaxis,” Cell, 109(5), pp. 611–623. Available at: 10.1016/S0092-8674(02)00755-9.

14. Garcia, G.L. and Parent, C.A. (2008) “Signal relay during chemotaxis.,” Journal of microscopy, 231(3), pp. 529–534. Available at: 10.1111/j.1365-2818.2008.02066.x.

15. Gulluni, F. et al. (2019) “Class II PI3K Functions in Cell Biology and Disease,” Trends in Cell Biology, 29(4), pp. 339–359. Available at: 10.1016/j.tcb.2019.01.001.

16. Hashimura, H. et al. (2019) “Collective cell migration of Dictyostelium without cAMP oscillations at multicellular stages,” Communications Biology, 2(1), p. 34. Available at: 10.1038/s42003-018-0273-6.

17. Hawkins, Phillip T. and Stephens, L.R. (2016) “Emerging evidence of signalling roles for PI(3,4) *P* 2 in Class I and II PI3K-regulated pathways,” Biochemical Society Transactions, 44(1), pp. 307–314. Available at: 10.1042/BST20150248.

18. Hawkins, P. T. and Stephens, L.R. (2016) “Emerging evidence of signalling roles for PI(3,4)P2 in class I and II PI3K-regulated pathways,” Biochemical Society Transactions, 44, pp. 307–314.

19. Hoeller, O. et al. (2013) “Two distinct functions for PI3-kinases in macropinocytosis,” Journal of Cell Science, p. jcs.134015. Available at: 10.1242/jcs.134015.

20. Hoeller, O. and Kay, R.R. (2007) “Chemotaxis in the Absence of PIP3 Gradients,” Current Biology, 17(9), pp. 813–817. Available at: 10.1016/j.cub.2007.04.004.

21. Iijima, M. and Devreotes, P. (2002) “Tumor Suppressor PTEN Mediates Sensing of Chemoattractant Gradients,” Cell, 109(5), pp. 599–610. Available at: 10.1016/S0092-8674(02)00745-6.

22. Kimmel, A.R. and Parent, C.A. (2003) “The signal to move: D. discoideum go orienteering.,” Science (New York, N.Y.), 300(5625), pp. 1525–1527. Available at: 10.1126/science.1085439.

23. Kriebel, P.W. et al. (2018) “Extracellular vesicles direct migration by synthesizing and releasing chemotactic signals,” Journal of Cell Biology, 217(8), pp. 2891–2910. Available at: 10.1083/jcb.201710170.

24. Kriebel, P.W., Barr, V.A. and Parent, C.A. (2003) “Adenylyl Cyclase Localization Regulates Streaming during Chemotaxis,” Cell, 112(4), pp. 549–560. Available at: 10.1016/S0092-8674(03)00081-3.

25. Lam, P. et al. (2012) “The SH2-domain-containing inositol 5-phosphatase (SHIP) limits neutrophil motility and wound recruitment in zebrafish,” Journal of Cell Science, p. jcs.106625. Available at: 10.1242/jcs.106625.

26. Li, X. et al. (2018) “Mutually inhibitory Ras-PI(3,4)P(2) feedback loops mediate cell migration.,” Proceedings of the National Academy of Sciences of the United States of America, 115(39), pp. E9125–E9134. Available at: 10.1073/pnas.1809039115.

27. Li, X. et al. (2021) “Reverse fountain flow of phosphatidylinositol-3,4-bisphosphate polarizes migrating cells,” The EMBO Journal, 40(4), p. e105094. Available at: 10.15252/embj.2020105094.

28. Loomis, W.F. (2014) “Cell signaling during development of Dictyostelium,” Developmental Biology, 391(1), pp. 1–16. Available at: 10.1016/j.ydbio.2014.04.001.

29. Loovers, H.M. (2007) “Regulation of phagocytosis in Dictyostelium by the inositol 5-phosphatase OCRL homolog Dd5P4,” Traffic, 8, pp. 618–628.

30. Nishio, M. (2007) “Control of cell polarity and motility by the PtdIns(3,4,5)P3 phosphatase SHIP1,” Nature Cell Biology, 9, pp. 36–44.

31. Pal, D.S. et al. (2019) “The excitable signal transduction networks: movers and shapers of eukaryotic cell migration.,” The International journal of developmental biology, 63(8-9–10), pp. 407–416. Available at: 10.1387/ijdb.190265pd.

32. Pirola, L. et al. (2001) “Activation Loop Sequences Confer Substrate Specificity to Phosphoinositide 3-Kinase α (PI3Kα),” Journal of Biological Chemistry, 276(24), pp. 21544–21554. Available at: 10.1074/jbc.M011330200.

33. Posor, Y. et al. (2013) “Spatiotemporal control of endocytosis by phosphatidylinositol-3,4-bisphosphate,” Nature, 499(7457), pp. 233–237. Available at: 10.1038/nature12360.

34. Postma, M. et al. (2004) “Sensitization of Dictyostelium chemotaxis by phosphoinositide-3-kinase-mediated self-organizing signalling patches,” Journal of Cell Science, 117(14), pp. 2925–2935. Available at: 10.1242/jcs.01143.

35. Ridley, A.J. et al. (2003) “Cell Migration: Integrating Signals from Front to Back,” Science, 302(5651), pp. 1704–1709. Available at: 10.1126/science.1092053.

36. Subramanian, B.C., Majumdar, R. and Parent, C.A. (2017) “The role of the LTB 4 -BLT1 axis in chemotactic gradient sensing and directed leukocyte migration,” Seminars in Immunology, 33, pp. 16–29. Available at: 10.1016/j.smim.2017.07.002.

37. Swaney, K.F. et al. (2015) “Novel protein Callipygian defines the back of migrating cells.,” Proceedings of the National Academy of Sciences of the United States of America, 112(29), pp. E3845–3854. Available at: 10.1073/pnas.1509098112.

38. Swaney, K.F., Huang, C.-H. and Devreotes, P.N. (2010) “Eukaryotic chemotaxis: a network of signaling pathways controls motility, directional sensing, and polarity,” Annual Review of Biophysics, 39, pp. 265–289. Available at: 10.1146/annurev.biophys.093008.131228.

39. Wallroth, A. and Haucke, V. (2018) “Phosphoinositide conversion in endocytosis and the endolysosomal system,” Journal of Biological Chemistry, 293, pp. 1526–1535.

40. Weijer, C.J. (2004) “Dictyostelium morphogenesis,” Current Opinion in Genetics & Development, 14(4), pp. 392–398. Available at: 10.1016/j.gde.2004.06.006.

41. Yamada, K.M. and Sixt, M. (2019) “Mechanisms of 3D cell migration,” Nature Reviews Molecular Cell Biology, 20(12), pp. 738–752. Available at: 10.1038/s41580-019-0172-9.

42. Zhou, K. et al. (1995) “A phosphatidylinositol (PI) kinase gene family in Dictyostelium discoideum: biological roles of putative mammalian p110 and yeast Vps34p PI 3-kinase homologs during growth and development.,” Molecular and Cellular Biology, 15(10), pp. 5645–5656. Available at: 10.1128/mcb.15.10.5645.

